# Reorganization of budding yeast cytoplasm upon energy depletion

**DOI:** 10.1101/468454

**Authors:** Guendalina Marini, Elisabeth Nüske, Weihua Leng, Simon Alberti, Gaia Pigino

**Affiliations:** Max Planck Institute of Molecular Cell Biology and Genetics, Dresden, Germany

## Abstract

Yeast cells, when exposed to stress, can enter a protective state in which cell division, growth and metabolism are downregulated. They remain viable in this state until nutrients become available again. How cells enter this protective survival state and what happens at a cellular and subcellular level is largely unknown. In this study, we used electron tomography to investigate the stress-induced ultrastructural changes in the cytoplasm of yeast cells. After ATP depletion, we observed a significant cytosolic compaction and an extensive cytoplasmic reorganization, as well as the emergence of distinct membrane-bound and membrane-less organelles. By using correlative light and electron microscopy (CLEM), we further demonstrate that one of these membrane-less organelles is generated by the reversible polymerization into large bundles of filaments of the eukaryotic translation initiation factor 2B (eIF2B), an essential enzyme in the initiation of protein synthesis. The changes we observe are part of a stress-induced survival strategy, allowing yeast cells to save energy, protect proteins from degradation, and inhibit protein functionality by forming assemblies of said proteins.

## Introduction

To survive in a constantly changing world, cells require mechanisms to cope with environmental fluctuations. For instance, the baker’s yeast *Saccharomyces cerevisiae* responds to sudden changes in nutrient abundance by adjusting its metabolism and growth rate. Under starvation conditions, yeast proliferation is efficiently shut down in favor of an energy-saving state that allows the cells to remain viable on slow metabolic activity (Werner-Washburne *et al.*, 1993; Choder, 1993; Ireland et al, 1994; Gray *et al.*, 2004). However, the structural changes that happen in the cells upon entry in this energy-saving state remain unclear.

The organization of the yeast cytoplasm is very dynamic and changes continuously, not only following the different stages of the cell cycle, but also in response to stress conditions such as heat shock, osmotic stress, and nutrient starvation (Meaden et al, 1999; Winderickx *et al.*, 2003; Petrovska *et al.*, 2014; Mourão *et al.*, 2014; Munna *et al.*, 2015; Joyner *et al.*, 2016; Munder *et al.*, 2016). The transition from a growing to a dormant state, induced through a severe lack of nutrients, is coupled to various physio-chemical changes, such as a lowered cytosolic pH, reduced cell volume and decreased molecule mobility in the cytoplasm (Ashe *et al.*, 2000; Joyner *et al.* 2016, Munder *et al.*, 2016), as well as physiological changes, such as a reduction in protein synthesis and in enzymatic activities (Werner-Washburne *et al.*, 1993; Gray *et al.*, 2004). Since the majority of metabolic reactions and signaling processes take place in the cytoplasm, stress-induced changes in its physical and chemical properties might be required for the cell to transition into a survival state. However, how the cytoplasm reorganizes under sudden stress conditions is largely unclear and comprehensive structural investigations of cytoplasmic changes occurring are missing.

Live-cell imaging has shown that several fluorescently tagged enzymes form assemblies in the cytoplasm of dormant cells (Narayanaswamy *et al.*, 2009; Noree *et al.*, 2010; Liu, 2010; Petrovska *et al.*, 2014; Riback *et al.*, 2017; Prouteau *et al.*, 2017). Electron microscopy imaging of three of these metabolic enzymes, cytidine triphosphate synthetase (CtpS, Barry *et al.*, 2014), glutamine synthetase (Gln1, Petrovska *et al.*, 2014) and target of rapamycin complex 1 (TORC1, Prouteau *et al.*, 2017), revealed that they assemble into filaments and that the filaments organize in bundles, visible in fluorescence microscopy. In all three cases, the polymerization process is reversible, and it is coupled to the inactivation of the enzymatic activities. These findings suggest that this stress-induced polymerization could be a common mechanism to downregulate the activity of essential metabolic enzymes during adverse environmental conditions.

In this study, we used electron tomography (ET) to investigate the reorganization of the cytoplasm of energy-depleted yeast cells, and we used correlative light and electron microscopy (CLEM) to study the behavior of the essential enzyme eIF2B (eukaryotic translation Initiation Factor 2B), a guanine nucleotide exchange factor and key initiator of protein synthesis. We show that energy-depleted yeast cells undergo a dramatic reorganization of the cytoplasm that involves the formation of distinct membrane-bound and membrane-less organelles, along with an almost two-fold increase in macromolecular crowding. Our results show that eIF2B decamers self-assemble into ordered bundles of filaments. We propose that eIF2B compartmentalization upon energy depletion could be a mechanism to store and protect the enzyme from denaturation, disassembly or vacuolar degradation, and could possibly contribute to the downregulation of its enzymatic activity. These results are consistent with a model in which cytoplasmic reorganization and enzyme regulation via self-assembly is an important survival strategy that enable cells to cope with extreme environmental conditions and stress.

## Results

### Energy depletion is accompanied by a reorganization of storage and membrane lipids

To investigate how the cytoplasm of energy-depleted yeast cells undergoes structural rearrangements, we have been mimicking sudden glucose starvation by feeding yeast cells with a non-hydrolyzable analogue of glucose, 2-deoxyglucose, and by blocking mitochondrial respiration with antimycin drug, *de facto* depriving cells of any possible source of ATP. After 15 minutes of treatment, we cryo-fixed the cells by high-pressure freezing to ensure the rapid immobilization of all macromolecular components in the cytoplasm, thus avoiding potential structural alterations that might occur during the slower process of standard chemical fixation (Frank, 2006). Subsequently, we imaged 70 nm to 150 nm thick sections of control and energy depleted yeast cells by dual-axis electron tomography and reconstructed the tomographic volumes from the aligned tilted images. By comparing the ultrastructure of log-phase growing cells with energy depleted cells, we found pronounced structural modifications between the two conditions (Figure 1; Video 1 for control; Video 2 and 3 for energy depletion), as energy depletion was characterized by extensive rearrangements of many cytoplasmic components, in particular lipids. We observed an increased number of small cytoplasmic lipid droplets (LDs), recognizable by their amorphous non-electron-dense content. They appeared fragmented and irregular (Figure 1, B and C; Video 2 and 3) compared to the fewer and larger LDs visible in control cells (Figure 1A; Video 1). The average LDs diameter in control cells was ~160 nm, with a measured area per vesicle of ~1.7 μm^2^. In energy-depleted cells, the diameter of lipid droplets was reduced to ~120 nm and the average area per vesicle to 0.9 μm^2^.

**Figure 1.**
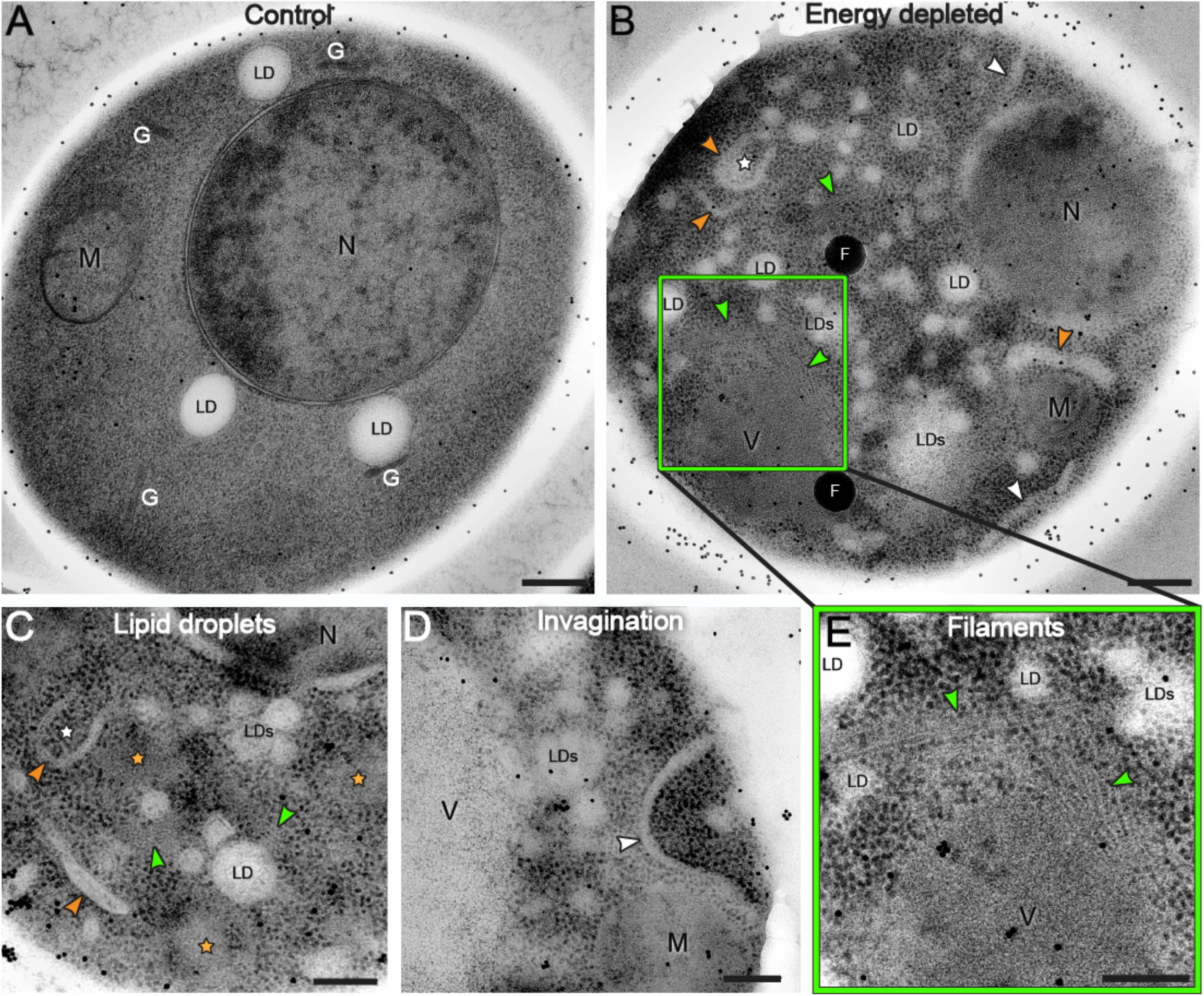
TEM images reveal cytoplasm reorganization in dormant yeast cells. Log-phase growing yeast cell (A) is compared to energy depleted yeast cells (B-D) showing that the cytoplasm undergoes a drastic reorganization upon sudden energy depletion. The vacuole is not visible in panel (A) because not included in this section of the cell. The disperse black dots in the images are gold beads of 15 nm in diameter used as alignment fiducials during tomography reconstruction. (B-D) Energy depleted cells contain numerous fragmented lipid droplets (LD and LDs, smaller droplets are not labeled), nascent autophagosomes (white stars in B and C), and elongated membranous structures (orange arrows). Ribosomes appear more densely packed in energy depleted cells compared to control cell (A), and areas of ribosome exclusion are also visible (orange stars in C). (B,C), elongated invagination of the cell membrane (white arrows) (B,D), and filamentous structures are visible in several parts of the cell (green arrows) (B,C, inset E). N = nucleus, M = mitochondria, G = Golgi, LD(s) = lipid droplet(s), V = vacuole, F = fluorescent beads. Scale bars: A and B = 300 nm; C-E = 200 nm.

Membrane lipids also underwent a great rearrangement through the accumulation of elongated membranous structures and the appearance of deep tonoplast invaginations in the cytoplasm (Figure 1, B and D; Video 2 and 3). The invaginations extended to 1 μm in length, with diameters ranging from 20 to 100 nm.

Interestingly, in energy-depleted cells we also observed double-layered membrane vesicles containing mostly ribosomes (Figure 1, B and C; Figure 4, B-D; Supplemental Figure S1 and Video 3). These resembled autophagosome vesicles, sometimes captured in an open, prefusion conformation (Figure 1, B and C). These structures could indicate early steps of the autophagy process, a process that is constitutive in cells, and can also be induced by stress (Noda and Ohsumi, 1998; Jin and Klionsky, 2014).

### Ribosome density and distribution change in energy-deprived cells

A previous study of energy-depleted yeast reported an increased mechanical stability of the cytoplasm and a ~7% reduction in cell volume (Munder *et al.*, 2016). Based on these observations, it was proposed that macromolecular crowding increases in stressed yeast cells. However, a direct quantification of cytoplasm density was never obtained. Since ribosomes take up a significant fraction of the cellular interior (Duncan and Hershey, 1983; Warner 1999), an accurate quantification of ribosome density can be used to account for changes in cytoplasmic crowding. In order to use ribosome density to quantify macromolecular crowding, we first verified that ribosome number does not change significantly between log-phase growing and dormant (energy-depleted) cells. Immunoblot analysis on ribosomal proteins of both 40S and 60S subunits (RPS4a and RPL32, respectively) showed no significant change in the quantity of ribosomes between untreated and energy depleted yeast cells (Figure 2E).

**Figure 2.**
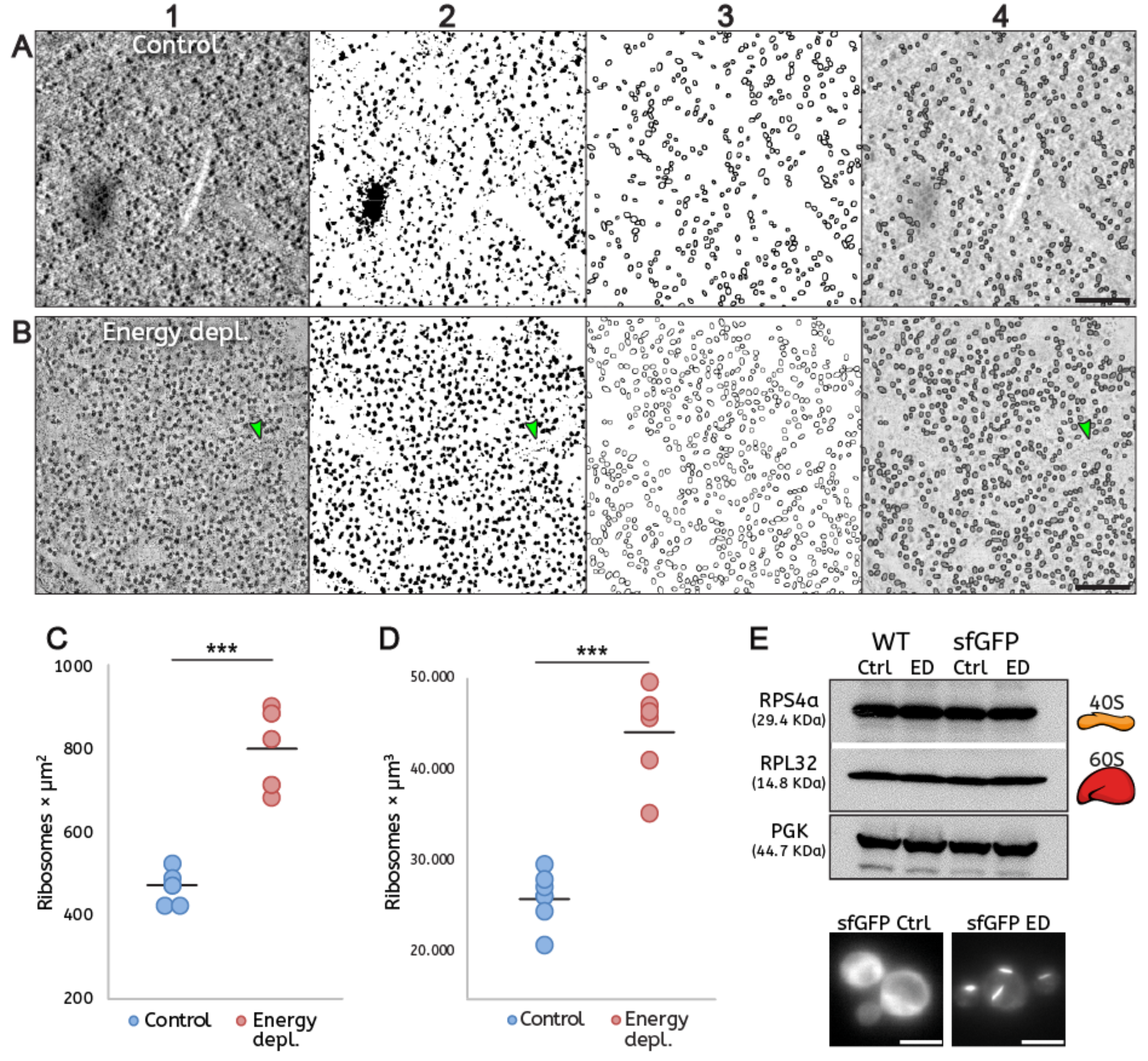
Quantification of ribosome density demonstrates macromolecular crowding in energy-depleted cells. (A-B) Automated quantification of ribosomes involving sequential (1) filtering, (2) binarization, (3) segmentation, and (4) particle detection on single tomographic slices of cells in (A) control conditions and (B) after energy depletion. Green arrows in panels B indicate a short filament. (C and D) The ribosome number is increased almost two-fold in the energy-depleted cells. (C) Ribosome density measured in tomographic slices. Five tomograms were analyzed per condition. ***P-value = 0.0001. (D) Ribosomes density is measured from tomographic volumes and assessed by manual counting. Six tomograms were analyzed per condition. ***P-value = 0.0001. (E) Western blot shows no significant change in the amount of RPS4a (40S) and RPL32 (60S) ribosomal subunits between log-phase growing (Ctrl) and energy-depleted (ED) cells in both wild-type and sfGFP-tagged eIF2B strains. Cells expressing sfGFP-tagged eIF2B were imaged immediately before Western blot analysis to validate the log-phase growth: diffuse GFP signal in the control condition (sfGFP Ctrl) and condensed GFP signal in energy depletion (sfGFP ED). Scale bars: A-B = 200 nm; E = 5 μm.

In our TEM tomograms, ribosomes appeared as extremely electron dense and were clearly distinguishable as dark dots in the cytoplasm, due to the presence of negatively charged rRNAs that are efficiently stained by contrast reagents used for sample preparation (Figure 2, A1 and B1; Supplemental Figure S2, A and B). To quantify the ribosome density, we took advantage of their sharp contrast and we developed a FIJI macro to automatically detect and count them in tomographic slices (Schindelin *et al.*, 2012). The images were processed by filtering, binarization, segmentation and particle detection (Figure 2, A and B). The accuracy of the automated quantification workflow was tested by comparison to ground truth data generated by manual counting of the ribosomes in single tomographic slices (Supplemental Figure S2). Ribosomes were manually counted in five cytoplasmic squares in every tomogram, avoiding areas that contained mainly large organelles such as vacuole, nucleus or mitochondria. Both quantifications showed that ribosome density increased almost twofold in energy-depleted cells (Figure 2, C and D; Supplemental Figure S2, C).

### Membrane-less compartments in energy-depleted cells have two distinct morphologies

Recent studies have proposed macromolecular crowding as one of the possible physical conditions to promote condensation of fluorescently labeled enzymes in the cytoplasm of energy depleted yeast cells (Petrovska *et al.*, 2014; Joyner *et al.* 2016; Munder *et al.*, 2016). We observed that, along with the dramatic increase in ribosome density, energy-depleted cells in our study showed also cytoplasmic areas from where ribosomes and larger macromolecular complexes were clearly excluded. These areas were never observed in control cells and showed a neat separation from the rest of the cytoplasm without being delimited by membranes. In particular, two main morphologies could be distinguished: some areas appeared as amorphous aggregate-like bodies (Figure 1C; Supplemental Figure S4, C – orange stars), other areas appeared as ordered self-assembly structures mostly composed of filaments organized in bundles (Figure 1, B and E; Figure 3C, F-H – green arrows; Figure 4 – white arrows; Videos 2 and 3 – white arrows). Both type of compartments did not show any obvious specific distribution in the cytoplasm, nor specific interaction with organelles or cellular components. The EM data suggest that these membrane-free areas of ribosome exclusion could represent specialized compartments for specific components of the cytoplasm, such as proteins linked to RNA and/or other enzymes, a possible mechanism of stress-induced segregation of such molecular components.

**Figure 3.**
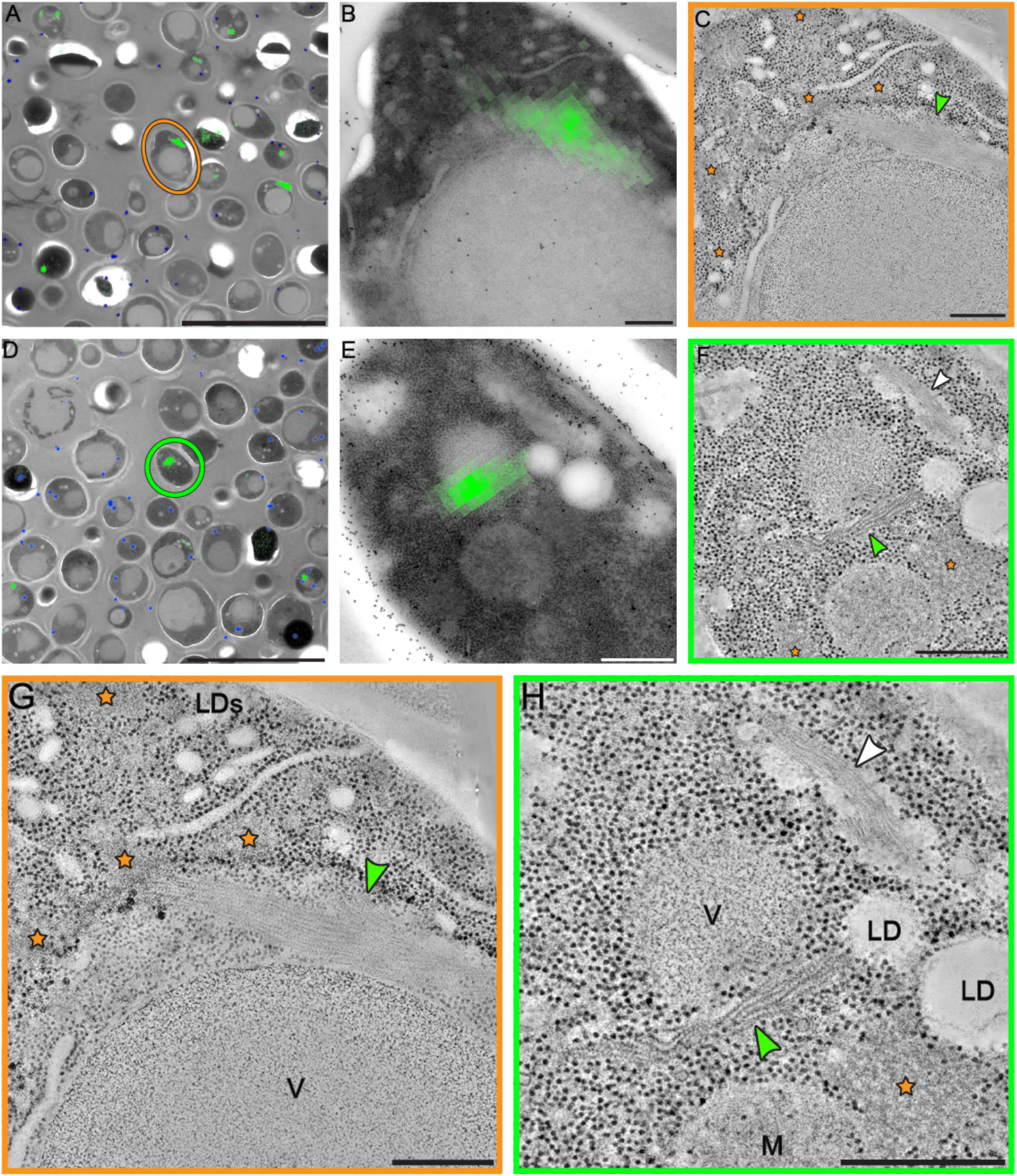
Correlative fluorescence and electron microscopy (CLEM) analysis reveals the organization of sfGFP-tagged eIF2B into bundles of parallel filaments. (A and D) The fluorescent signal of sfGFP-eIF2B is overlaid on low-magnification TEM images of energy-depleted yeast cells previously embedded in Lowicryl HM-20 and then sectioned. The orange circle in (A) and green circle in (D) highlight cells selected for further tomographic analysis. (B and E) Close ups of the cells highlighted in A and D, respectively. (C,F-H) Slices through tomographic reconstructions. (G and H) Magnified views of the tomographic slices shown in C and F, respectively. Bundles of filamentous structures (green arrows) corresponding to the fluorescence signal. eIF2B organizes in ordered, membrane-less arrays of filaments. Energy-depleted cells also contain other non-membrane bound compartments: some have an amorphous appearance (orange stars), whereas others comprise filamentous structures (H) (white arrow). These filaments, which do not display a fluorescent signal, have a different morphology to the eIF2B filaments and their protein content is unknown. LD(s) = lipid droplet(s); M = mitochondrion; V = vacuole. Scale bars: A, D = 10 μm; B-C, G = 200 nm; E-F, H = 500 nm.

**Figure 4.**
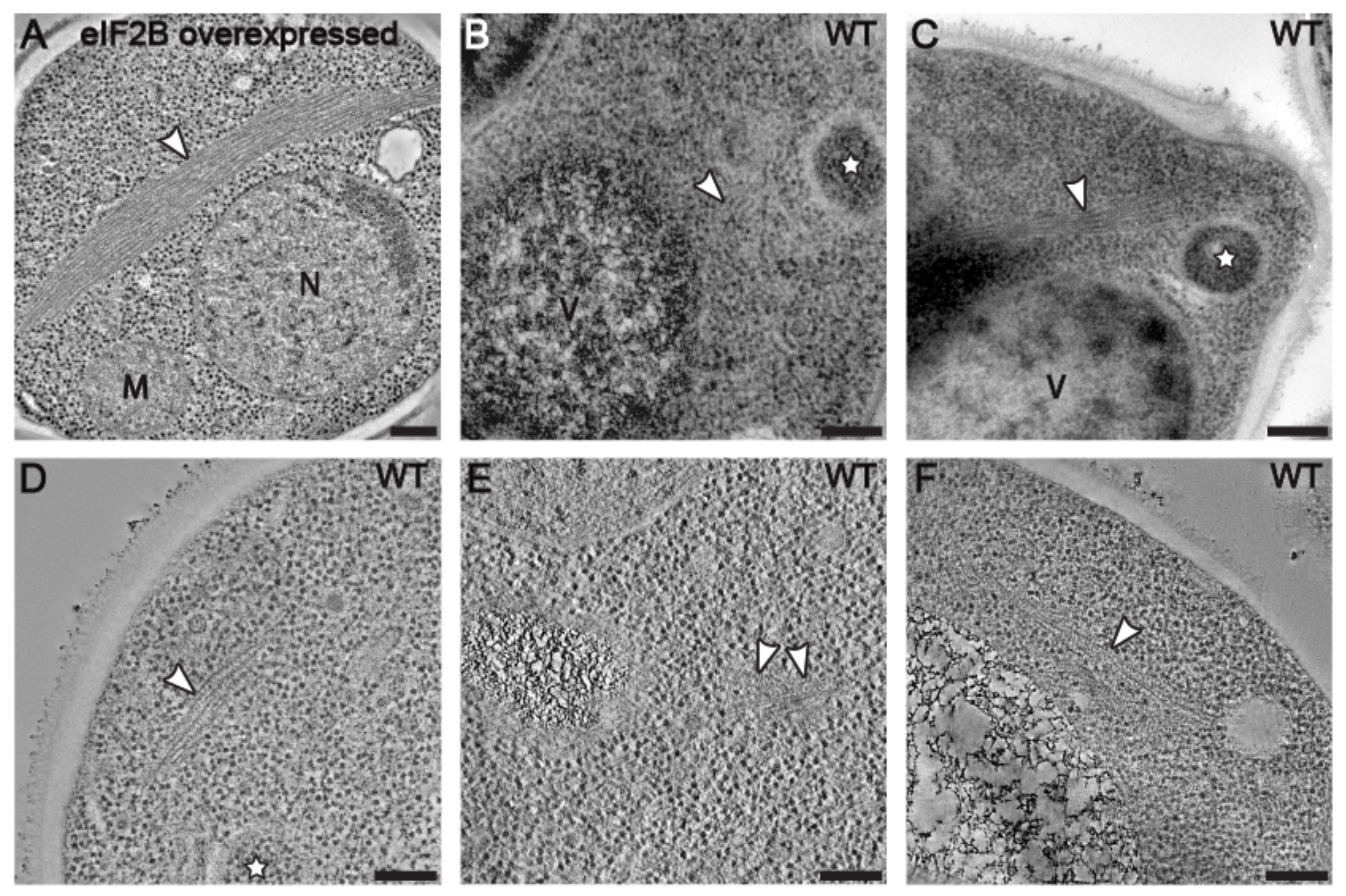
Filament formation is independent from sfGFP tagging and eIF2B expression levels. Average images of 20 tomographic slices of wild-type yeast cells showing (A) overexpressed eIF2B forming large bundles of filament in the cytoplasm, and (B-F) endogenously expressed eIF2B forming smaller bundles of filaments. (B-D) White stars label autophagosomes, recognized as double membrane vesicles containing ribosomes. Scale bars = 200 nm.

### eIF2B organize rapidly into highly ordered compartments upon energy depletion

Our TEM observations are in agreement with fluorescence microscopy studies reporting that a large number of enzymes, including eIF2B, are able to form assemblies upon nutrient/energy deprivation (Campbell *et al.*, 2005; Narayanaswamy *et al.*, 2009; Noree *et al.*, 2010; Liu, 2010). However, so far only Gln1, CtpS and TORC1 enzymes have been shown to self-assemble into highly ordered polymers in such stressful condition by TEM (Petrovska *et al.*, 2014; Barry *et al.*, 2014; Prouteau *et al.*, 2017). It is not known whether other enzymes assemble into polymers and whether this is a common occurrence in the cytoplasm of energy-depleted cells.

The eIF2B translation initiation factor is a key guanine nucleotide exchange factor highly conserved throughout eukaryotes that plays a fundamental role in the initiation of protein expression (Kurat *et al.*, 2006; Hinnebusch and Lorsch, 2012; Jennings and Pavitt 2014; Gordiyenko *et al.*, 2014 and 2019). It is known to form elongated and reversible fluorescent assemblies in yeast and mammalian cells under different stress conditions (Campbell *et al.*, 2005; Noree *et al.*, 2010; Nüske *et al.*, 2018; Hodgson *et al.*, 2019). Therefore, we tested whether eIF2B could be a constituent of one of the highly ordered membrane-less compartments we described above. To test this hypothesis, we performed CLEM on energy-depleted yeast cells overexpressing a sfGFP-tagged version of the eukaryotic translation initiation factor 2B (eIF2B) enzyme. Firstly, we followed the sfGFP-eIF2B signal in live-cell fluorescence microscopy (Supplemental Figure S3 – Video 4). In agreement with a previous study (Noree *et al.*, 2010), eIF2B enzymes showed a diffuse cytoplasmic signal in log-phase growing cells (Supplemental Figure S3 A and E). About 15 minutes after energy depletion, most of eIF2B molecules were concentrated into foci and elongated structures, and cell growth and division were heavily slowed down, if not arrested (Supplemental Figure S3, B and C). This condition lasted until replenishment of ATP, when the fluorescence signal promptly diffused again, and cells quickly re-entered the cell cycle (Supplemental Figure S3, D and E). The reversibility of the condensation process has also been described for self-assemblies made by CtpS (Barry *et al.*, 2014), Gln1 (Petrovska *et al.* 2014) and TORC1 (Prouteau *et al.*, 2017), with clear regulatory functions for all the three enzymes.

After imaging the fluorescent signal, we prepared yeast cells overexpressing sfGFP-eIF2B, both untreated and energy-depleted cells, for electron microscopy imaging. Then we correlated the fluorescence signals of the condensed eIF2B with the TEM images and located cell sections containing the labeled structures of our interest (Figure 3; Supplemental Figure S4). On those sections, we acquired dual-axis electron tomograms (Mastronarde 1997). The tomograms of energy-depleted cells showed that eIF2B polymerizes into a specific type of membrane-less cytoplasmic compartment that is composed of long zigzagged filaments packed in big bundles (Figure 3C, F-H). The morphology of eIF2B filaments and bundles was distinct from the morphology of other non-membrane bound compartments of stressed cells (Figure 3, F and H – green and white arrows, respectively). A single energy-depleted cell could contain several eIF2B bundles, with overexpressing cells showing larger bundles (Supplemental Figure S4). eIF2B filaments were never observed in untreated cells from the same strain and cell culture (Video 1).

In all analyzed tomograms of stressed cells, macromolecular complexes in the size range of ribosomes were excluded from the space occupied by the eIF2B bundles. Additionally, eIF2B filaments and bundles were never observed to be associated with autophagosomes, which often engulfed cytoplasmic components in energy-depleted cells (Figure 1, B and C; Figure 4, B-D; Supplemental Figure S1 – white stars; Video 3). These data indicate that compartmentalization of eIF2B in the bundles could be a mechanism to escape autophagy and protect the enzyme from vacuolar degradation.

### sfGFP-tag does not interfere with eIF2B polymerization

It has been reported that fluorescent proteins have a natural dimerizing affinity and tend to form aggregates or even higher-order structures (Zacharias *et al.*, 2002; Snapp *et al.*, 2003). This might pose a problem for applications in which fluorescent proteins are used to visualize the cellular localization, dynamics, and oligomeric state of a protein. Here, we used three distinct approaches to test whether eIF2B self-assembly happens intrinsically or might be caused or enhanced by the GFP tag: i) we used the non-dimerizing version of GFP (sfGFP, Costantini *et al.*, 2012); ii) we imaged eIF2B by immunofluorescence using the short HA polypeptide chain as tag; iii) we acquired electron tomograms of the native untagged eIF2B in energy-depleted wild-type yeast.

For the immunofluorescence experiments, we labeled with HA-tag two distinct subunits of the eIF2B complex, the regulatory subunit Gcn3 (α) and the catalytic subunit Gcd1 (γ), and we stained the samples with antibodies against the HA tag (Supplemental Figure S5). The analysis confirmed that eIF2B formed assemblies upon energy depletion also when a different tag than sfGFP was used.

TEM analysis of energy-depleted wild-type cells using untagged eIF2B, either endogenously expressed or overexpressed, revealed bundles of filaments showing the same morphology observed in CLEM experiments (Figure 4). As observed for sfGFP-tagged cells, bundles of untagged eIF2B were larger in overexpressing strains (Figure 4A) than in wild type cells (Figure 4, B-F). These results demonstrate that the formation of eIF2B filaments and bundles is an intrinsic property of the enzyme and that the sfGFP tag is not involved in this process. Additionally, we observed that the overexpression of eIF2B does not affect the mechanism of filament formation but leads only to bulkier bundles.

### eIF2B filaments comprise repeats of decameric units

To investigate the 3D organization of the filaments in the bundle, we measured the filaments length and the distance between filaments from the raw tomograms (Figure 5 – Video 5). The filaments length varied from ~45 nm to ~850 nm. The shorter filaments comprise three or four copies of eIF2B decamers and were usually observed at the periphery of the bundle. Filaments were mostly aligned in regular rows in the bundle, with a between-row spacing of ~13 nm and within-row spacing of ~26 nm (Figure 5C; Figure 6A). Diameter, periodicity and spacing of eIF2B filaments in the bundle differed from the ones of previously characterized enzymatic polymers, including those made of other metabolic enzymes such as CtpS, Gln1 and TORC1 (Barry *et al.*, 2014; Petrovska *et al.*, 2014; Prouteau *et al.*, 2017). In stressed cells, we also observed filaments with a similar morphology and subunit repeat (~6 nm) to the ones described for Gln1 (Petrovska *et al.*, 2014), with smaller diameter, shorter between-filament spacing, and smoother longitudinal striations than eIF2B filaments (Figure 3, F and H – white arrows).

**Figure 5.**
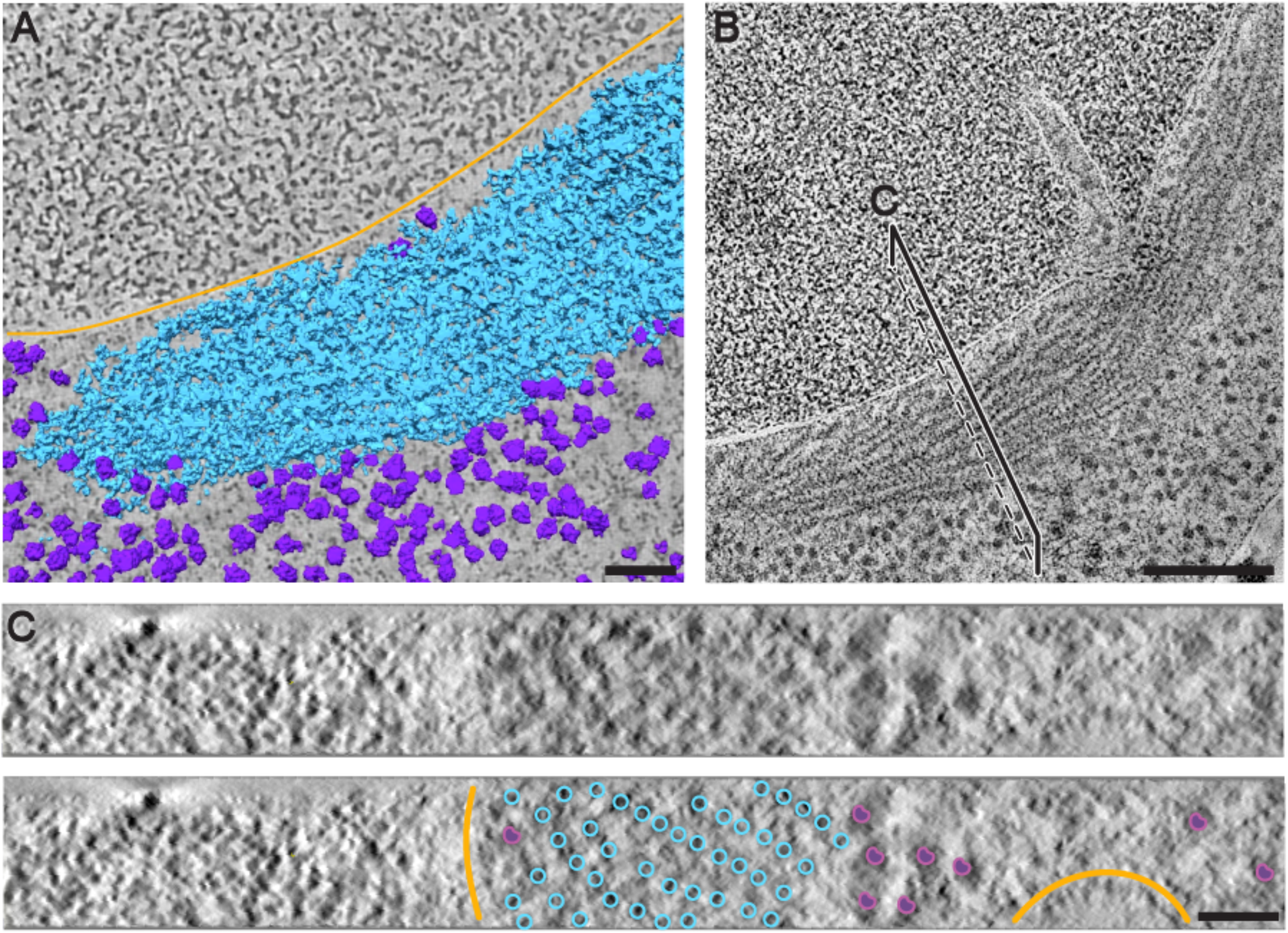
Segmentation and cross-sectional view of a eIF2B bundle showing the ordered arrangement of eIF2B filaments. (A) Segmentation of membranes (orange), ribosomes (purple) and eIF2B filaments (cyan) of a cell overexpressing sfGFP-tagged eIF2B. (B) Tomographic slice through a larger field of view of the same tomogram. eIF2B filaments are densely packed and parallel to each other in the bundle. (B) The black line indicates a cutting plane through the bundle corresponding to the cross-sectional view in (C). (C) eIF2B filaments follow a regular pattern with a center-to-center inter-filament spacing of approximately 20 nm. Positions of the filaments in the cross section have been annotated according to the segmentation. Scale bars: A = 50 nm; B = 200 nm; C = 40 nm.

**Figure 6.**
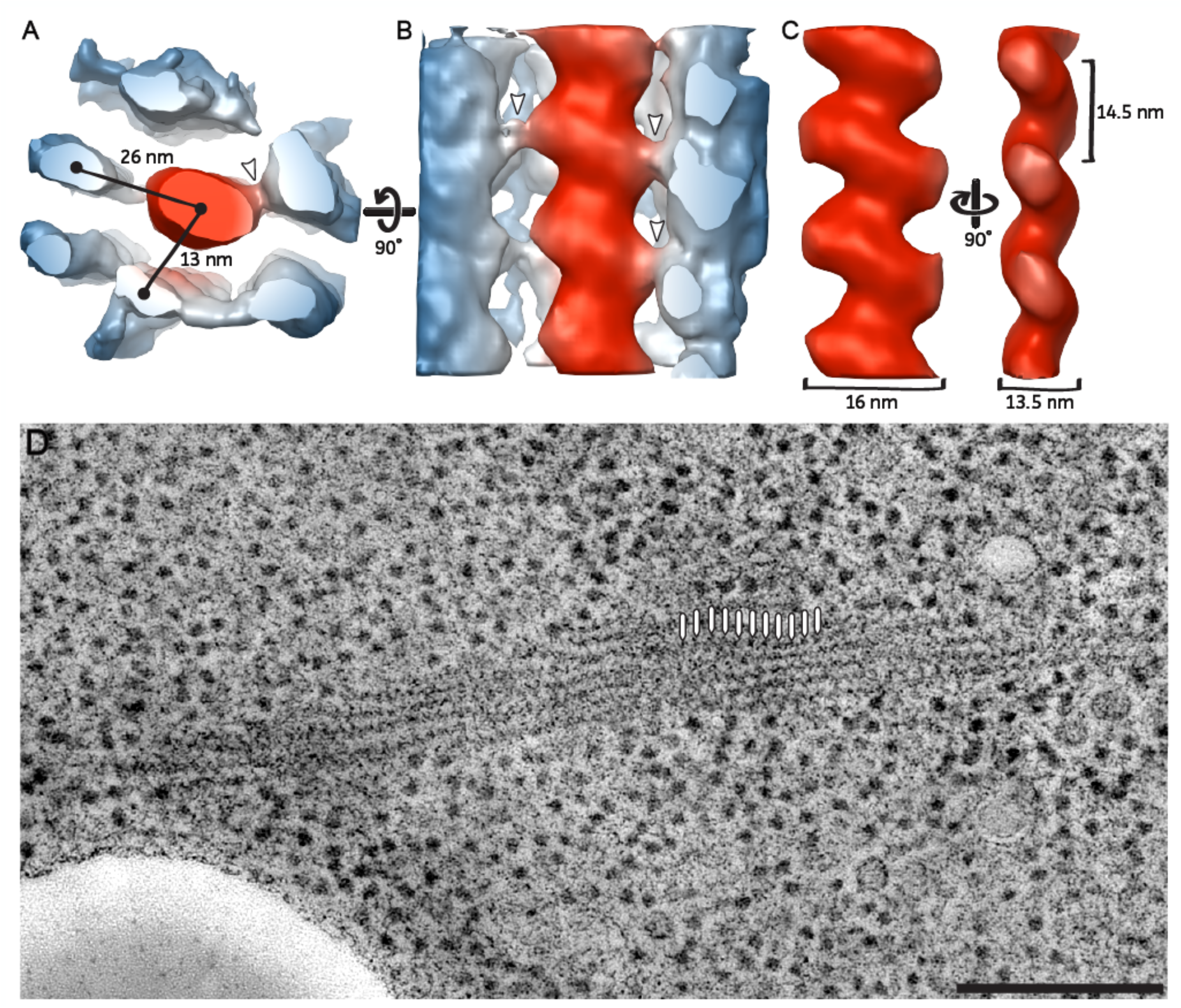
3D organization of parallel filaments in the eIF2B bundle. Subtomogram averaging of three repeating eIF2B units along a filament. (A-B) Neighboring filaments (in blue) are still visible in the average, which is indicative of their roughly consistent position around a central filament (in red). The center-to-center distance between the central filament and the surrounding ones is ~26 nm for filaments in the same row and ~13 nm for filaments in parallel rows. Lateral connections are visible between the central filament and the surrounding ones (white arrows). (B-C) The longitudinal view of the central filaments has a zigzag shaped structure. The central filament shows a 14.5 nm repeat, a long diameter of 16 nm and a short diameter of 13.5 nm. (D) Average of 10 tomographic slices showing the eIF2B bundle used for the 3D model reconstruction of the filaments and the periodicity at which particles have been picked (white lines). Scale bars: A-C = 10 nm; D = 200 nm.

Because of their apparent repeating structure, eIF2B filaments could represent polymers of a single protein complex. We used a subtomogram averaging approach to obtain a 3D model of the repeating unit. The 3D model showed a zigzag pattern with a 14.5 nm repeat (Figure 6C) in agreement with the shape of the pattern visible in all the raw tomograms. A single filament measured ~16 nm for the long diameter and 13.5 nm for the short diameter (Figure 6C). Contact points between central and neighboring filaments were visible in raw tomograms and in the 3D model (Figure 6, A and B – white arrows) as they protruded from both sides of the zigzag shape. The regularity of the inter-filament packing, where a central filament is surrounded by six or seven equidistant neighboring ones (Figure 5; Figure 6A), suggested that the observed lateral connections could contribute to keeping distance between molecules and a regular pattern of filaments in the bundle.

Although the sample preparation protocol used for ET in this study does not allow for the generation of high resolution 3D models by subtomogram averaging, we extracted the volume of the eIF2B decamer from the cryoEM map of eIF2B/eIF2αP complex (Adomavicius *et al.*, 2019; Gordiyenko *et al.*, 2019), and we fitted a gaussian filtered version into our 3D model of the filament. The fitting showed that periodicity and curvatures of three decameric eIF2B units were well accommodated in our 3D model (Figure 7). The eIF2B decamers needed to be rotated of ~45° from their vertical axis to fit in the zigzag pattern of the filament, thus the only point of contact between units appears to be through Gcd6 ε-subunits.

**Figure 7.**
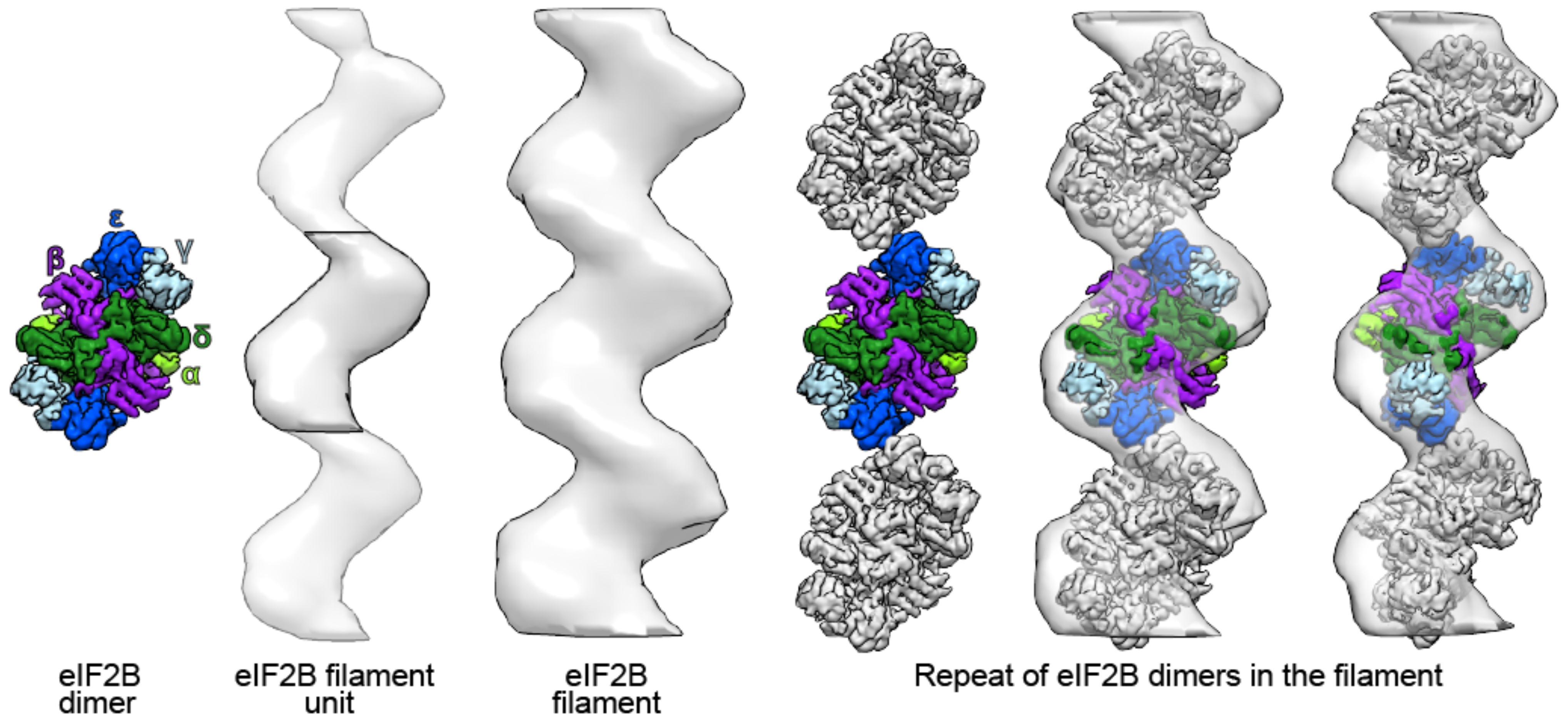
Proposed organization of eIF2B decamers in the filament. The repeating units in eIF2B filaments have a size and overall shape similar to the Gaussian filtered maps of the cryo-EM eIF2B decamer [Adomavicius et al., 2019; Gordiyenko et al., 2019]. The fitting of three density maps within three periodic eIF2B units in the 3D model of the filament obtained by subtomogram averaging suggests a possible stacking of eIF2B decamers in the polymerized form.

Since it is known from previous work that eIF2 and eIF2B colocalizes to eIF2-eIF2B bodies in fluorescent assays of *S. cerevisiae*, *C. albicans*, and human astrocytic cells (Pavitt *et al.*, 1998; Campbell *et al.*, 2005; Egbe *et al.*, 2015; Taylor *et al.*, 2010; Noree *et al.*, 2010; Hodgson *et al.*, 2019), we fitted the full eIF2(αP)/eIF2B complex (Adomavicius *et al.*, 2019; Gordiyenko *et al.*, 2019) into the eIF2B filament model in order to understand whether eIF2 can also be present in the polymer. In this case the β and γ subunits of eIF2 protruded outside the filament. However, subunit eIF2α was mostly included in the filament model and the position of the eIF2 protrusions corresponded to the positions of the lateral connections we observe as contact points between the filaments. The eIF2 arm appeared to be very flexible and only transiently in contact with eIF2B, explaining why in our 3D model of the filament we observe frequent lateral connections but never with a regular repetition.

## Discussion

In this paper, we demonstrate that entrance of yeast cells into a stress survival state is associated with a broad reorganization of molecules and the formation of structures in the cytoplasm. We reveal that the reorganization involves a massive rearrangement of membranes and lipids as well as the formation of compartments that are not surrounded by membranes.

### Lipid droplets change their morphology upon energy depletion

The observed changes in lipid droplet (LDs) morphology might be indicative of the dramatic effects that the scarcity of nutrients/energy has on the regulation of lipid metabolism (Rußmayer *et al.*, 2015). In yeast cells, LDs are particularly dynamic throughout the growth phase and depending on the nutritional status of the cell (Kurat *et al.*, 2006; Bozaquel-Morais *et al.*, 2010). Cells that progressively reach the stationary phase and experience gradual nutrient depletion, are known to accumulate and enlarge LDs, due to their shift from a lipolytic to a lipogenic metabolism, allowing cells to produce basal levels of energy from fatty acids for cellular maintenance (Madeira *et al.*, 2015). Upon replenishment of glucose, large amounts of sterols and fatty acids are mobilized and, as a result, LDs are decreased in number and size (Kurat *et al.*, 2006).

In our experiments, yeast cells were grown until early-mid log-phase and then exposed to an acute energy starvation, without previous accumulation of LDs. Since starvation is known to switch yeast cell metabolism towards β-oxidation of fatty acids, which yields more energy per gram than carbohydrates like glucose (Gray *et al.* 2004; Kurat *et al.*, 2006), LDs present in the cell are strongly mobilized and appeared smaller and fragmented, as a possible consequence of an activation of lipid metabolism. In addition, LDs fragmentation could contribute to the increase in total surface area accessible to lipases, enhancing lipolytic activity and contributing to the minimal production of energy during stress conditions, as it has been proposed in animal cells (Ariotti *et al.*, 2012; Hashimoto *et al.*, 2012; Thiam and Beller, 2017).

### Plasma membrane invaginations accumulate as a consequence of cell shrinkage

The pronounced plasma membrane invaginations present in energy-depleted cells were morphologically similar to those observed in yeast cells under severe hyperosmotic stress. This stress condition is usually associated with dehydration and rapid shrinkage of the cell volume (Morris *et al.*, 1986; Blomberg and Adler, 1992; Gervais and Marechal, 1994; Martinez de Maranon *et al.*, 1996; Slaninova et al, 2000; Simonin *et al.*, 2007; Dupont *et al.*, 2010). Concomitantly, due to the poor compressibility of biological membranes (Thiery *et al.*, 1967), cell shrinkage is associated with wrinkling of the tonoplast (Adya *et al.*, 2006). Therefore, the formation of plasma membrane invaginations that we observed in energy-depleted cells can well be the consequence of the reduction in cell volume, attributed to a sudden water loss that follows the acute energy depletion in *S. cerevisiae* (Munder *et al.*, 2016).

Moreover, upon energy replenishment, the cell surface needs to rapidly increase, to allow cells to return to their normal volume. Such a rapid expansion of the plasma membrane requires ‘membrane reservoirs’ that provide surface area and buffer membrane tension. We speculate that the accentuated depth of plasma membrane invaginations in starved cells allows the rapid shrinkage of the membrane into reservoirs, which then provide additional surface for prompt re-expansion of the plasma membrane upon energy replenishment.

### Ribosome density is a measure of increased macromolecular crowding

Munder *et al.* (2016) measured a dramatic decrease in particle mobility in the cytoplasm of energy depleted cells, which was proposed to result from increased molecular crowding and condensation of the cytoplasm. However, the measured ~7% reduction in the cell volume would hardly be sufficient to induce this pronounced effect on particle mobility. Therefore, we directly quantified ribosomes in 3D electron tomograms to verify and measure changes in molecular crowding between control and energy depleted yeast cells. While the total ribosome number remained unchanged between the two conditions, we observed an almost two-fold increase in ribosome density in energy depleted cells. Based on these data, we estimate a theoretical cell volume reduction of about 42%, which is far more pronounced than that measured by Munder *et al.* (2016). This discrepancy could be explained by the concomitant enlargement of the vacuole, which has previously been reported to occur in response to starvation (Desfougères *et al.*, 2016; Joyner *et al.*, 2016) and also during the progression of autophagy (Noda and Ohsumi, 1998). Indeed, our quantification is in line with previously measured cytoplasmic volume reduction of 30% in yeast cells upon glucose starvation (Joyner *et al.*, 2016). Therefore, we propose that a combination of cell volume reduction and vacuole enlargement is most likely responsible for the increase in molecular crowding observed in starved yeast cells. Regardless of the exact sequence of events causing the changes in the physical-chemical properties of the cytoplasm, our analysis provides a direct quantification of the contribution of molecular crowding in yeast cells.

### Increased macromolecular crowding is associated with assembly and compartment formation

Changes in molecular crowding are thought to have profound effects on cytoplasm organization and on cell physiology. *In vitro* experiments show that the assembly of individual enzymatic species is triggered by the addition of crowding agents (Petrovska *et al.*, 2014; Woodruff *et al.*, 2017). Consequently, crowding has been proposed to play an important role in the formation and stabilization of assemblies and membrane-less compartments in cells. The increased crowding of stressed yeast cells that we report here could indeed represent one of the physical parameters that promote the formation of protein-specific assemblies, such as the bundles of eIF2B filaments. The formation of eIF2B filaments requires the diffusion of soluble components and the incorporation of these components into a nucleation center. Filament nucleation and growth could be promoted by the increased crowding conditions in the cytoplasm, because the resulting higher protein concentration favors molecular interactions and chemical reactions (Rivas *et al.*, 2001; Zhou *et al.*, 2008). Conversely, excessive crowding could decrease molecular motion, especially of large particles, resulting in diminished protein assembly and binding and consequently a reduced rate of chemical reactions (Trappe *et al.*, 2001; Miermont *et al.*, 2013; Joyner *et al.*, 2016; Munder *et al.*, 2016). This suggests that filament nucleation and elongation must occur when crowding does not entirely prevent molecular mobility. Since we observed maximal crowding already after 15 minutes of energy depletion (Supplemental Figure S3), the recruitment of molecules into filaments must happen very rapidly at the beginning of this process. Hence, we propose that an increased macromolecular crowding in the cell favors and stabilizes assemblies and membrane-less compartments, but it does not necessarily cause nor initiate their formation in the cytoplasm.

There is increasing evidence that a sudden intracellular acidification, induced by energy starvation, triggers condensation of several enzymes in yeast cells (Petrovska *et al.*, 2014, Munder *et al.*, 2016, Rabouille and Alberti, 2017). This pH drop is thought to change surface charges of molecules, producing new repulsive and attractive molecular interactions in the cytoplasm. Therefore, pH change in the cytoplasm may be a key initiator of enzyme-specific assemblies. Even though the chemical nature of the signal that triggers assembly of eIF2B and other enzymes in energy depleted cells needs to be further elucidated, the process must occur under very low energy levels and before the cytoplasm becomes too crowded. Thus, stressed cells transition from a stage in which cellular components are allowed to dynamically reorganize, to a stage where the cytoplasm is condensed and rather static, as reflected in the reduced mobility of organelles and foreign tracer particles (Munder *et al.* 2016).

### Filaments and bundles form mainly via polymerization of eIF2B decamers

The rapid organizing of eIF2B in highly ordered bundles of filaments in the cytoplasm of energy depleted cells suggests that eIF2B filament formation is a specific adaptation to conditions in which energy levels are low. Using immunolabeling and tomography on WT cells, we were able to exclude that filament formation is triggered or affected by sfGFP-tagging, therefore confirming that filaments and bundles formation is an intrinsic property of eIF2B. It has been shown that energy depleted wild type yeast cells undergo translational arrest (Ashe *et al.*, 2000; Nüske *et al.*, 2018), whereas eIF2B mutated strains with a reduced ability of forming filaments keep on translating proteins longer after energy depletion (Nüske *et al.*, 2018). These findings seem to indicate that eIF2B filament formation promotes downregulation of translation and suggests that eIF2B activity is inhibited upon energy depletion. This suggests that filament formation could be a mechanism to silence eIF2B enzymatic activity in cells undergoing energy depletion.

Comparison of the eIF2B cryo-EM structure from *S. cerevisiae* (Adomavicius *et al.*, 2019; Gordiyenko *et al.*, 2019) with the periodicity and the size of the filament repeating unit, indicates that filaments are formed by polymerization of intact eIF2B decamers, rather than simply aggregation (Figure 7). More in detail, the fitting suggests that the longitudinal axis of eIF2B decamers might be rotated by ~45° in their assembled form, making them interact only through the catalytic Gcd6 ε-subunits (Figure 7). Gcd6 ε-subunit, together with Gcd1 γ-subunit, composes the “catalytic core” of eIF2B, which catalyzes the GDP/GTP exchange reaction (Williams *et al.*, 2000; Gomez and Pavitt, 2000). This proposed eIF2B arrangement in the filament would partially occlude the catalytic sites of both Gcd6 ε-subunit of eIF2B decamers, suggesting a possible mechanism for enzymatic inhibition.

Even though in fluorescent microscopy experiments eIF2B foci are known to colocalize with their reaction substrate, the eIF2 complex (Pavitt *et al.*, 1998; Campbell *et al.*, 2005; Noree *et al.*, 2010), we could not accommodate full densities of bound eIF2 molecules into our 3D model of the eIF2B filament (Adomavicius *et al.*, 2019; Gordiyenko *et al.*, 2019). Most of the filament density is occupied by eIF2B dimers, indicating that they are generated by polymerization of eIF2B complexes alone. Nevertheless, we cannot completely rule out the presence of eIF2 in the structure of the filament, as it might be only transiently bound to eIF2B filaments (Campbell *et al.*, 2005).

In summary, we have described the structural rearrangements taking place in the cytoplasm of energy depleted yeast cells (Figure 8). We propose that these changes allow cells to save energy and to protect essential cellular components, like eIF2B, from denaturation and/or vacuolar degradation. The reversible self-assembly of eIF2B, and potentially other enzymes, aimed for storage and inhibition of their energy demanding activities, could help cells to endure unfavorable environmental conditions. It will be interesting to investigate whether this cytoplasmic reorganization can be a general response for other proteins, in other species and to other kind of stresses.

**Figure 8.**
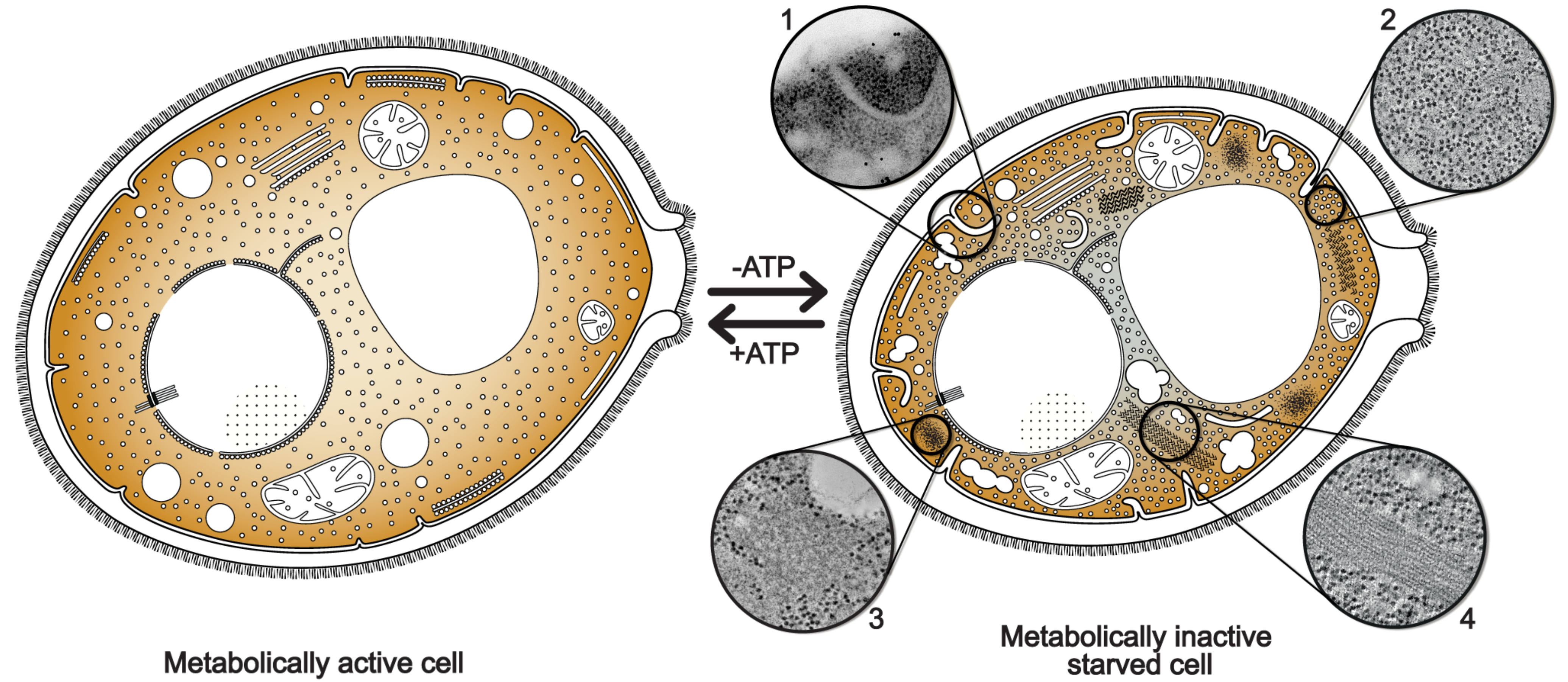
Energy depletion induces a massive reorganization of cytoplasmic structures in yeast cells. ATP-depleted cells undergo volume reduction, with consequent formation of plasma membrane invaginations (1) and a pronounced increase in macromolecular crowding (2). Numerous membrane-less compartments of amorphous (3) or highly ordered morphology, such as eIF2B filaments and bundles (4), appear in the cytoplasm. Energy-depleted cells are also characterized by the fragmentation of lipid droplets, suggesting a switch to β-oxidative metabolism of fatty acids. All these cellular rearrangements result in a “solidification” of the cytoplasm that could function as an energy saving state and as a protective mechanism for molecular components from denaturation and/or vacuolar degradation, to keep them readily available when favorable growth conditions are restored.

## Materials and methods

### Yeast strains, media and energy depletion

Wild type *Saccharomyces cerevisiae* W303 cells and the two strains overexpressing untagged and sfGFP-tagged eIF2B on the C-terminus of the Gcn3 α-subunit were grown in an orbital shaker (180 rpm) at 25°C in YPD medium containing 1% (w/v) yeast extract, 2% peptone, and 2% glucose. Detailed information on the generation of the mutant strains are published in a related study (Nüske *et al.*, 2018).

Stress treatment was carried out on cells grown to mid-log phase and cellular growth was monitored by optical density (OD) at 600 nm. To induce energy depletion (ATP depletion), cells at 0.5 OD_600nm_ were washed twice and then incubated in SD (synthetic dropout) complete medium containing 20 mM 2-deoxyglucose (2-DG; Carl Roth GmbH, Karlsruhe, Germany) and 10 M antimycin-A (Sigma-Aldrich, Steinheim, Germany) for at least 15 minutes. These two chemicals were used to block glycolysis and mitochondrial respiration, respectively. This treatment reduces intracellular ATP levels by more than 95% (Serrano, 1977). Yeast samples were then incubated in an orbital shaker at 30°C for 1 hour. Log phase yeast cells expressing only endogenous untagged eIF2B were treated in both complete and energy depleted medium in the same way as described above.

### Fluorescence microscopy

Yeast cells were imaged in concanavalin A-coated 4-well Matek dishes. Fluorescence microscopy of live (time-lapse movies) and fixed cells were acquired using a Deltavision microscope system with softWoRx 4.1.2 software (Applied Precision), 100× 1.4 NA UPlanSApo oil immersion objective, and CoolSnap HQ2 camera (Photometrics) at 1024×1024 (or 512×512) pixels. Exposure time was 0.3 seconds and the interval between consecutive frames in the time lapse was 15 minutes.

We determined the exact time required for yeast cells to enter the dormant state upon energy depletion by analysis of the fluorescence microscopy time-lapse movies. This information was required to select dormant cells for imaging in TEM and compare their ultrastructure with that of growing cells.

### Immunostaining

For immunostaining, cells expressing GCD1-γ-HA and Gcn3-α-HA were fixed via treatment with 3.7% paraformaldehyde (EMS, Hatfield, USA) for at least 30 minutes followed by 45 minutes incubation in spheroplasting buffer (100 mM phosphate buffer pH 7.5, 5 mM EDTA, 1.2 M Sorbitol (Sigma-Aldrich, Steinheim, Germany), Zymolyase (Zymo Research, USA) at 30°C with mild agitation. Spheroplasts were permeabilized with 1% triton X-100 (Serva, Heidelberg, Germany), washed and incubated with mouse anti-HA primary antibody (1:2000; Covance, USA) and goat anti-mouse-HRP (1:5000; Sigma, Saint Louis, USA).

### High pressure freezing and freeze substitution of yeast cells

Yeast cells, harvested by vacuum filtration, as described in Bertin and Nogales (2016), were transferred to 100 μm deep membrane carriers and high pressure frozen with a Leica EM PACT2 or Leica EM ICE freezers (Leica Microsystems, Wentzler, Germany). All samples were processed by freeze substitution in a Leica AFS2 temperature-controlling machine (Leica Microsystems, Wentzler, Germany) and then embedded in resin using two distinct protocols for untagged and sfGFP-tagged yeast strains.

#### Untagged eIF2B yeast cells

High pressure frozen samples were freeze substituted using 1% osmium tetroxide, 0.1% uranyl acetate (wt/vol) and 5% H_2_O (vol/vol) in glass distilled acetone. Freeze substitution was carried out at −90°C for 36 hours before raising the temperature steadily to −30°C in 4°C per hour. The samples were kept at −30°C for 5 hours before they were brought to 0°C in steps of 4°C per hour. Samples were washed with acetone and infiltrated with increasing concentrations (25, 50, 75 and 100%; 2 hours each) of EPON resin (Electron Microscopy Sciences). 100% EPON solution was exchanged two times in 12-hour steps. Resin infiltrated samples were then UV polymerized at 60°C for 48 hours. Samples were cut into 150 to 200 nm sections with an ultramicrotome (Ultracut UCT; Leica) and Ultra 35° diamond knife (Diatome). The sections were mounted on Formvar-coated slot grids (Science Services).

#### Yeast cells sfGFP-tagged on the C-terminal of the Gcn3 α-subunit of eIF2B

Vitrified yeast cells were freeze substituted with 0.1% (wt/vol) uranyl acetate and 4% (vol/vol) water in acetone at −90°C as described in Kukulski *et al.* (2011). The samples were then embedded in Lowicryl HM-20 (Polysciences Inc.) and cut into 70, 100 and 150 nm thick sections using an ultramicrotome (Ultracut UCT; Leica) with an Ultra 35° diamond knife (Diatome).

### Correlative light and electron microscopy

For CLEM imaging, sections of yeast cells expressing sfGFP-tagged Gcn3 (α)-subunit of eIF2B, embedded in Lowicryl HM-20 (Polysciences Inc.) (Kukulski *et al.*, 2011), were mounted on Formvar-coated finder grids (Science Services) and incubated with quenched blue FluoSphere® fiducials (Molecular Probes, 200 nm diameter, excitation/emission maxima at 365/415 nm) diluted 1:500 for 10 minutes in the dark. Grids were mounted on a glass slide with VectaShield (Vector Laboratories, Inc., Burlingame, USA) and imaged with a Zeiss Axioplan2 wide-field fluorescence CCD upright microscope to record the fluorescence signal in sections of the embedded yeast cells. Images were acquired with a 10x objective lens to generate a grid overview and a 100x objective lens (na. 1.4) to image the fluorescence in individual cells. Cells were imaged with a green GFP channel (488 nm) for the sfGFP-tagged eIF2B and a UV channel (nm) for the blue FluoSphere® fiducials. The correlation (overlay) between the images obtained in fluorescence microscopy and EM micrographs or tomographic slices was performed with AMIRA 2D-5D visualization and analysis software (ThermoFisher Scientific).

### Electron tomography

All sections from resin embedded samples were stained with 1% (wt/vol) uranyl acetate in 70% (wt/vol) methanol for 5 minutes and 0.4% lead citrate for 3 minutes. Colloidal gold particles (15 nm in diameter) were added to both surfaces of the sections to serve as fiducial markers for tilt series alignment. Tomographic series were acquired in dual-axis tilt scheme (±60° and 1° increments) with SerialEM (Mastronarde, 2005), using a FEI Tecnai F30 TEM (300 kV) equipped with Gatan US1000 CCD camera. Pixel size ranged between 7-12Å/px. Tomograms reconstruction was performed using the IMOD and ETomo packages (Mastronarde, 1997).

### Subtomogram averaging

Subtomogram averaging was done with PEET software from the IMOD package (Heumann *et al.*, 2011). Subtomograms of 54×54×54 voxels (40×40×40 nm) were automatically picked along eIF2B filaments every 15 nm. A final average was calculated from a total of ~500 particles. A loose mask was used to refine the central filament average.

Semi-automatic segmentation of structures of interest in the tomograms was obtained with the software SuRVoS (Luengo *et al.*, 2017). The USCF Chimera package was used for 3D visualization, rendering, and animation of the reconstructed volumes (Pettersen *et al.*, 2004). For eIF2B fitting, eIF2(αP)/eIF2B cryo-EM maps from Adomavicius *et al.*, 2019 (EMDB 4404) and from Gordiyenko *et al.*, 2019 (EMDB 4544) have been segmented based on eIF2B and eIF2 subunits from their corresponding atomic models (PDB 6I3M and 6QG1, respectively, whose modelling was based on the eIF2B crystal structure of *S. pombe* from Kashiwagi *et al.*, 2016). Surfaces of eIF2 subunits have been hidden and all eIF2B subunits have been gaussian filtered at 1.5 Å. The fitting in the 3D model of the filament has been performed using the “fit” command in UCSF Chimera (Pettersen *et al.*, 2004), treating all subunits as a part of a single model.

## Supporting information

3D tomographic reconstruction of a non-stressed yeast cell. Scale bar= 500 nm

Video2. 3D tomographic reconstruction of an energy-depleted yeast cell. eIF2B is unlabelled and overexpressed. Scale bar= 500nm

Video3. 3D tomographic reconstruction of an energy-depleted yeast cell. eIF2B is unlabelled and endogenously expressed. Scale bar= 500nm

Video4. The fluorescent signal of eIF2B is distributed differently between stressed yeast cells and log-phase growing cells. Scale bar = 10 microns

Video5. Segmentation of a bundle of eIF2B filaments in a reconstructed tomogram.

## Acknowledgments

We acknowledge Prof. JR. Warner and Ms. SV. Buhl, from the Albert Einstein College of Medicine (NY) for kindly donating the L3 (TCM) monoclonal and the L30/S4 polyclonal ribosome antibodies. We thank Prof. T. Müller-Reichert for sharing the Amira protocol of automated filament tracing, and D. Richter for cloning HA-tagged eIF2B yeast strains. We are grateful for the support of T. Furstenhaupt and K. Gibson, B. Borgonovo, A. Bogdanova in the EM and Protein Expression and Purification Facilities at the Max Planck Institute of Molecular Cell Biology and Genetics (MPI-CBG). We thank O Gonzalez for IT support, D Diener and F Jug for substantial advice and theoretical input. This work was supported by the Dresden International Graduate School for Biomedicine and Bioengineering (DIGS-BB), granted by the German Research Foundation (DFG) in the context of the Excellence Initiative. Simon Alberti acknowledges funding by the Volkswagen Life? Initiative and the Deutsche Forschungsgemeinschaft (AL 1061/5-1).

## Author contributions

GM: experimental design, data acquisition, data analysis and interpretation, manuscript writing. EN: experimental design, data acquisition, data interpretation. WL: support for data acquisition. SA: conception and design, manuscript writing. GP: conception and design, data analysis and interpretation, manuscript writing.

---

**Supplemental Figure S1.**
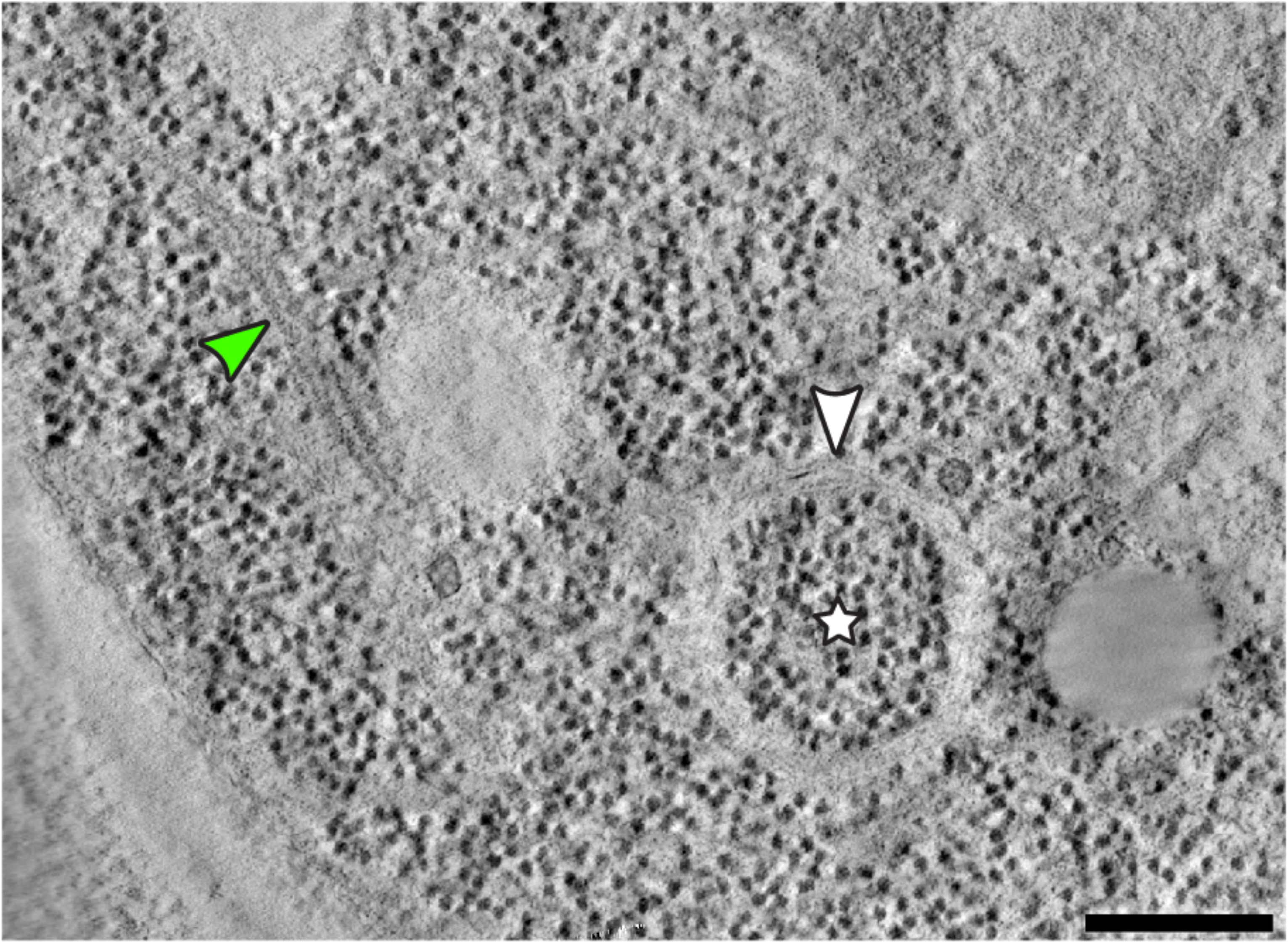
A double membrane vesicle (white arrow) is visible int the cytoplasm of an energy-depleted yeast cell. It contains tightly packed ribosomes in the lumen (white star) and it might be identified as an autophagosomal vesicle in the process of bringing its content to the vacuole for degradation. The green arrow points to a filament in the cytoplasm of the energy-depleted yeast cell. Scale bar = 200 nm.

**Supplemental Figure S2.**
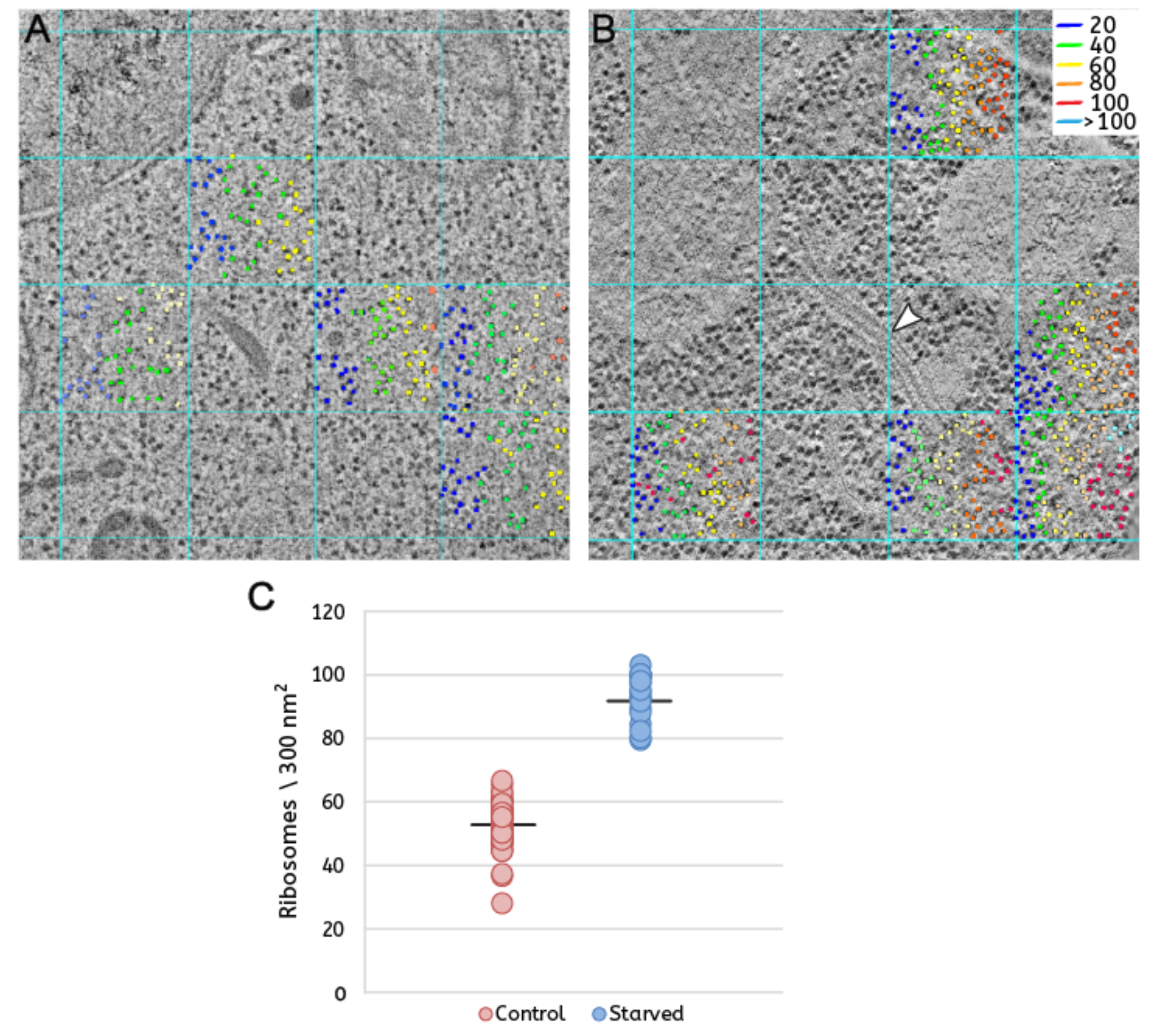
Manual method for ribosome counting. A 300 × 300 nm grid is superimposed on the central slice of a tomogram. Ribosomes in 5 cytoplasmic squares are counted, avoiding those comprising mainly large organelles, vacuole or nucleus. Ribosomes in the upper and left edge of each chosen square are included in the count. (A) Yeast control cells have usually 50-70 ribosomes per 300 nm^2^ area. (B) Energy depleted yeast cells have usually 100 or more ribosomes per 300 nm^2^ area. White arrow highlights a filament in the stressed cell. (C) The result of the manual ribosome counting is shown in the plot. Five tomograms are analyzed for each condition. The almost two-fold increase (42%) in ribosome density in the stressed yeast cells is confirmed.

**Supplemental Figure S3.**
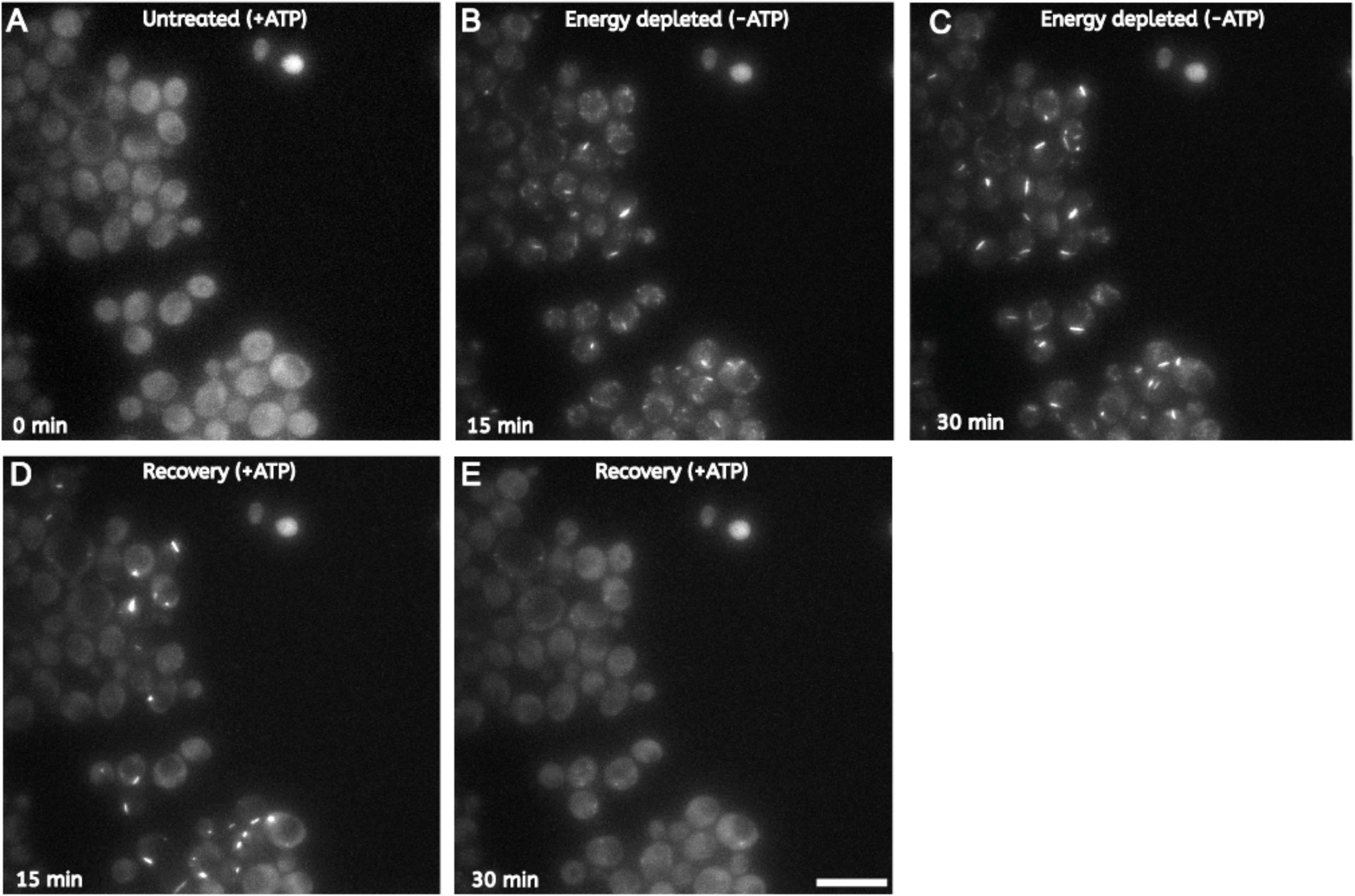
eIF2B enzymes change their cytoplasmic distribution between log-phase growing and energy-depleted yeast cells. (A) According to the fluorescent signal, the sfGFP-labeled eIF2B shows a diffuse distribution throughout the cytoplasm in log-phase growing and dividing cells. (B) Within 15 minutes from energy depletion, the fluorescence signal of sfGFP-tagged eIF2B is condensed into foci-like and elongated structures. (C) After 30 minutes of energy depletion, the number of foci is reduced in favor of larger condensates. (D, E) The fluorescent signal rapidly diffuses upon energy replenishment, and cells re-enter the cell cycle. Scale bar = 10 μm.

**Supplemental Figure S4.**
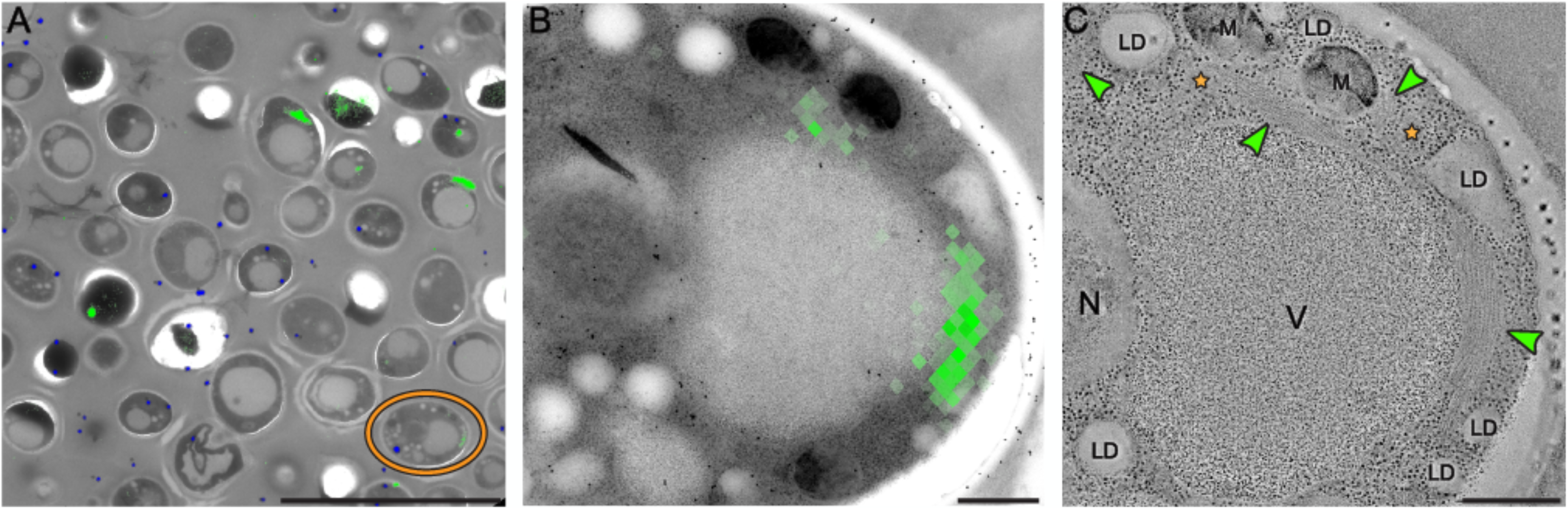
Correlative fluorescence and electron microscopy (CLEM) analysis reveals organization of sfGFP-tagged eIF2B into bundles of parallel filaments. (A) The fluorescent signal of sfGFP-eIF2B is overlaid on low-magnification TEM image of energy-depleted yeast cells embedded in Lowicryl HM-20 and sectioned. (B) Close up of the cell highlighted in A. (C) Magnified views of the tomographic slices shown in C. Two big bundles of filamentous structures and one small one (green arrows) corresponds to the fluorescence signal. Small non-membrane bound compartment with amorphous appearance are not fluorescent and labelled with orange stars. LD = lipid droplet; M = mitochondrion; V = vacuole; N = nucleus. Scalebars: A=10 μm, B and C=500 nm.

**Supplemental Figure S5.**
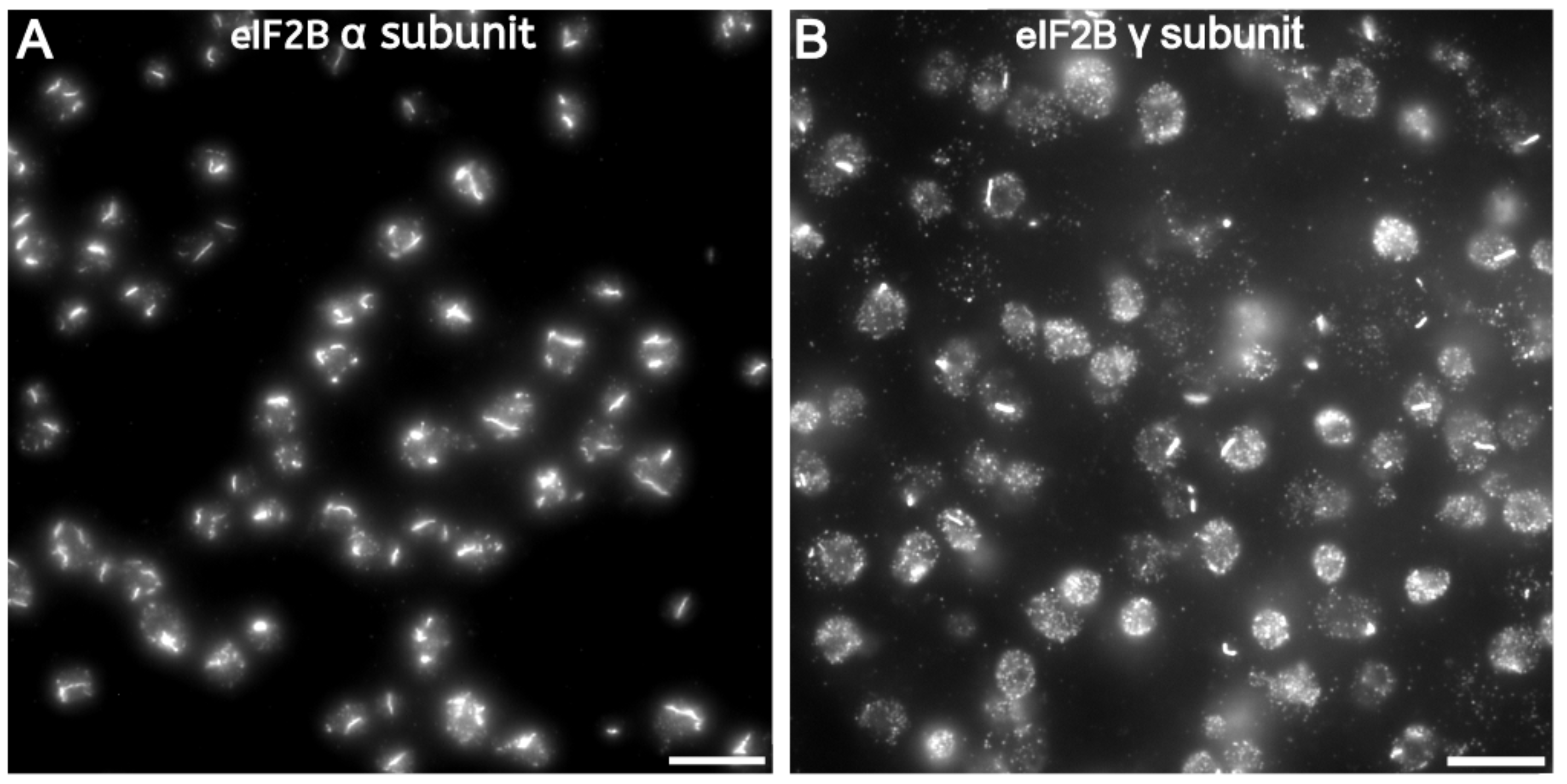
Immunofluorescence labeling of two HA-tagged eIF2B subunits show filament bundles in energy-depleted yeast cells. (A) Immunolabeling of the HA-tagged Gcn3 (α) and (B) Gcd1 (γ) subunits of eIF2B show that the formation of filament bundles in energy-depleted cells occurs in the absence of the sfGFP tag. This demonstrates that separation of eIF2B in non-membrane bound compartments is not induced or enhanced by the sfGFP tag. Scale bars = 10 μm.

**Video1.**
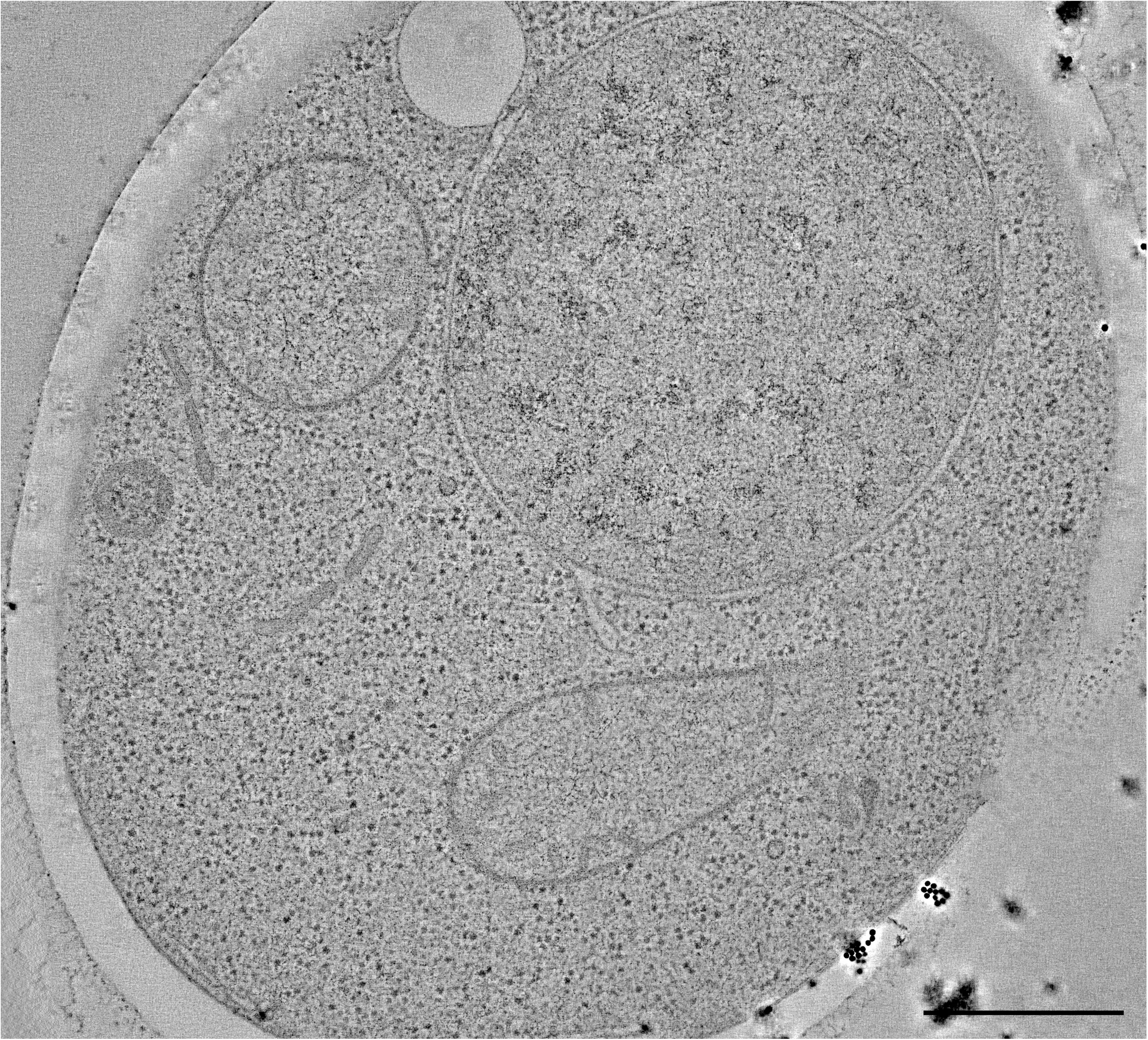
3D tomographic reconstruction of a non-stressed yeast cell. Scale bar= 500 nm

**Video2.**
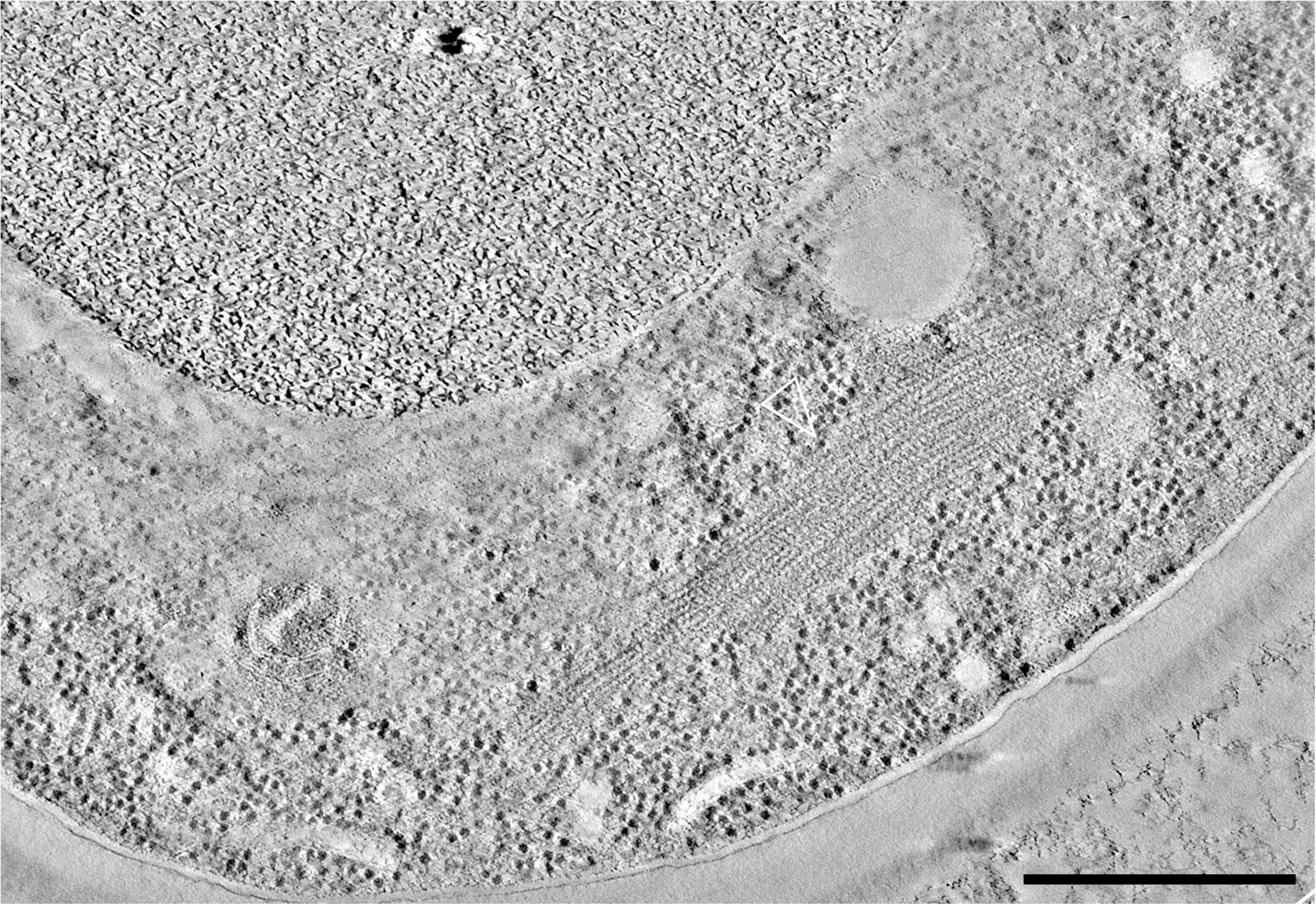
3D tomographic reconstruction of an energy-depleted yeast cell. eIF2B is unlabelled and overexpressed. Scale bar= 500nm

**Video3.**
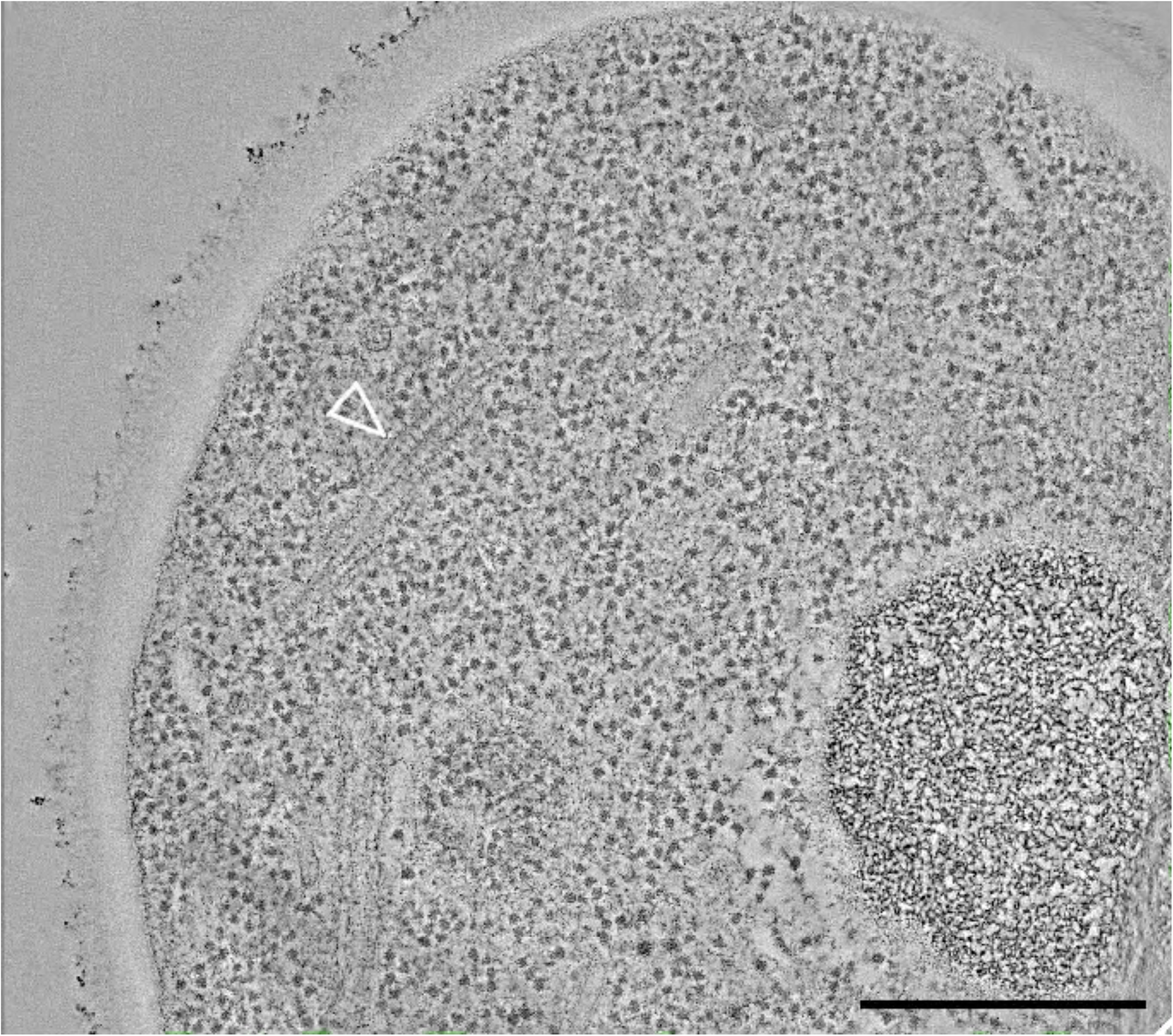
3D tomographic reconstruction of an energy-depleted yeast cell. eIF2B is unlabelled and endogenously expressed. Scale bar= 500nm

**Video4.**
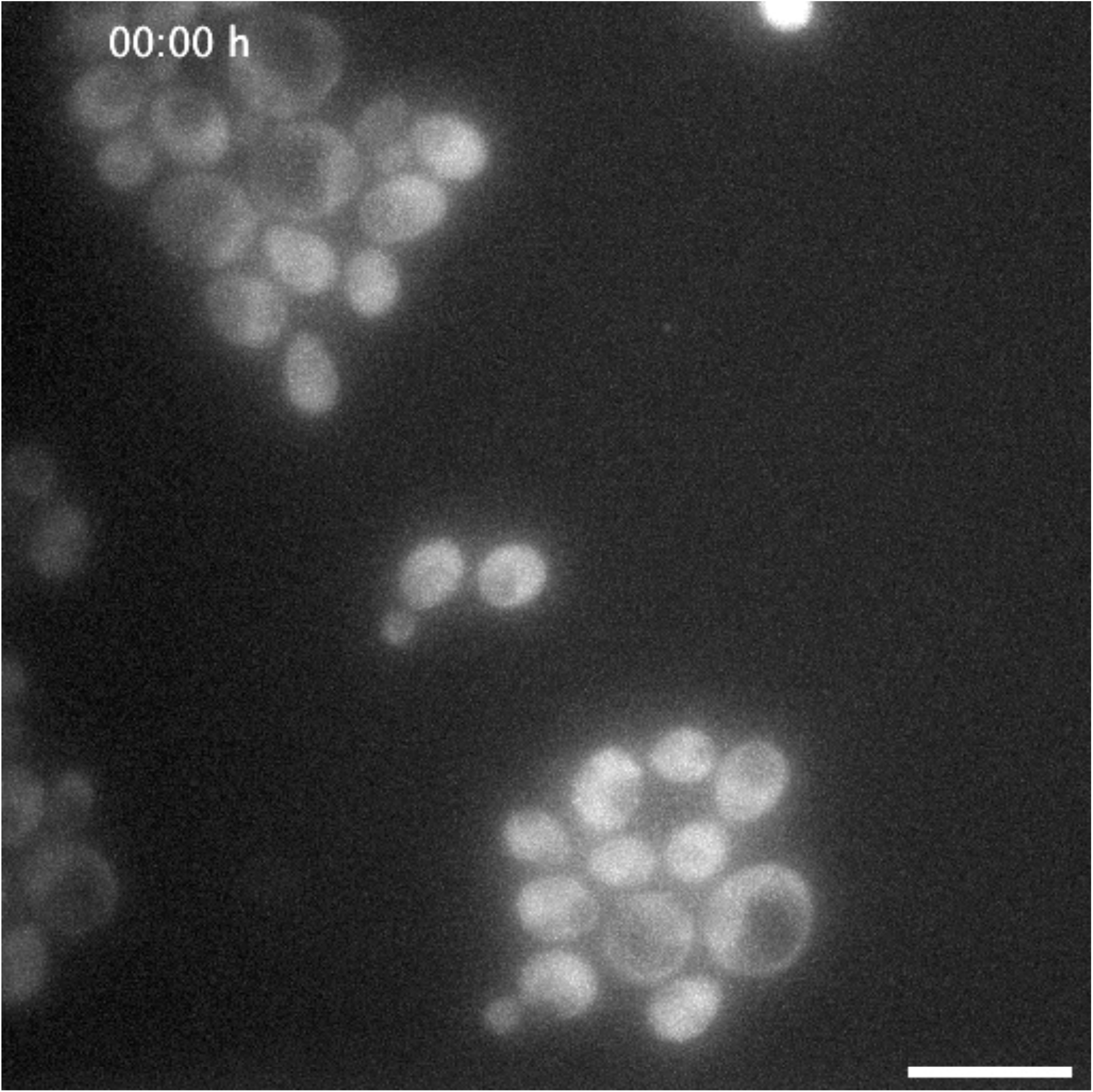
The fluorescent signal of eIF2B is distributed differently between stressed yeast cells and log-phase growing cells. According to the fluorescent signal, the sfGFP-labeled Gcn3 (α) subunit of eIF2B present a diffuse distribution throughout the cytoplasm in log-phase growing and dividing cells. On the contrary, the fluorescence is concentrated into condensed foci-like or elongated structures in stressed cells that have been energy depleted for 15 minutes at pH 5.5. Scale bar= 10 μm

**Video5.**
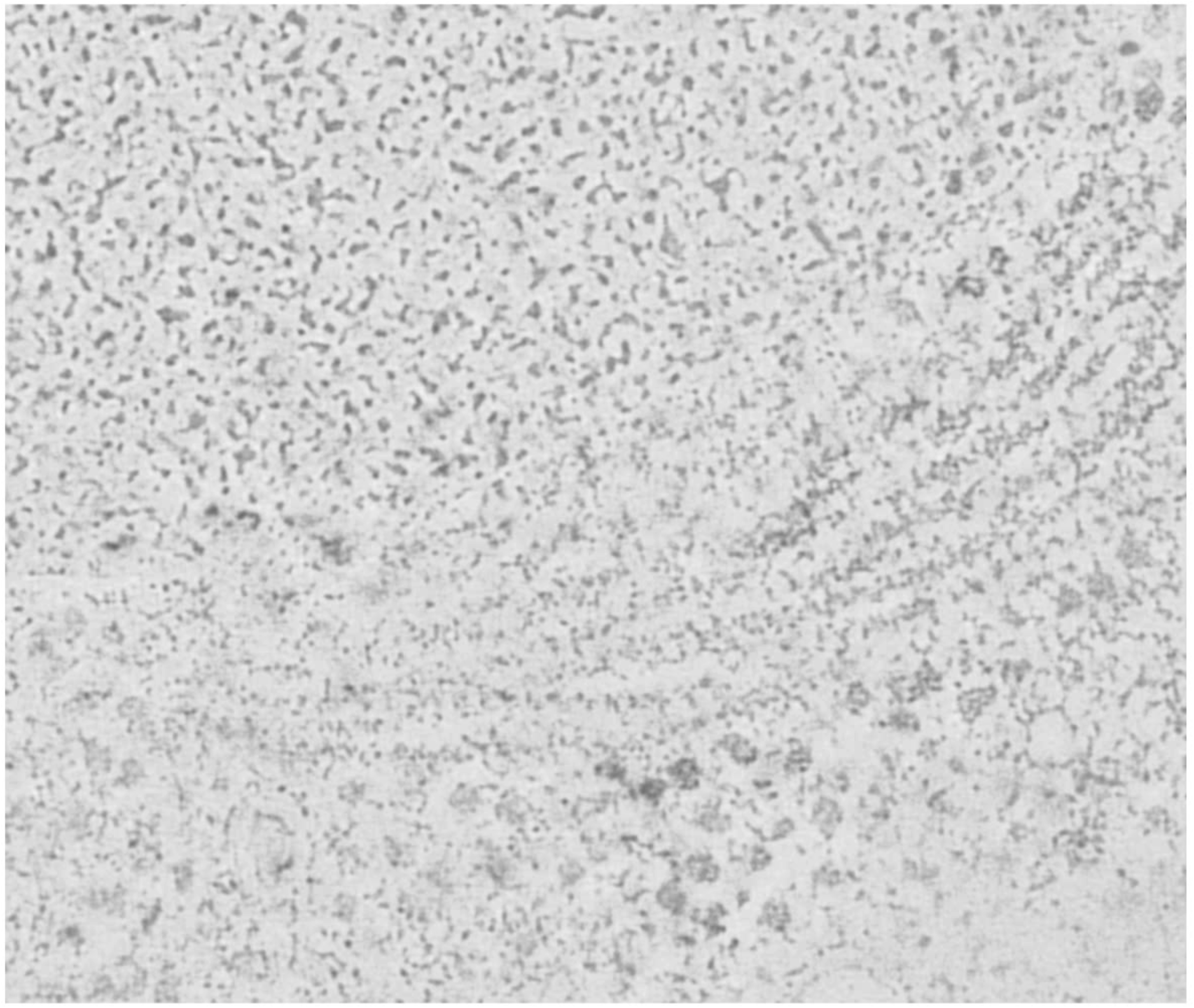
Segmentation of a bundle of eIF2B filaments in a reconstructed tomogram using Super Region Volume Segmentation software [Luengo et al., 2017]. Automated segmentation of ribosomes (purple) and eIF2B filaments (cyan) in the tomographic reconstruction of a cell overexpressing sfGFP-tagged eIF2B.

## REFERENCES

Adomavicius T, Guaita M, Zhou Y, Jennings MD, Latif Z, Roseman AM, Pavitt GD (2019). The structural basis of translational control by eIF2 phosphorylation. Nat Commun 10, 2136.

Adya AK, Canetta E, Walker GM (2006). Atomic force microscopic study of the influence of physical stresses on *Saccharomyces cerevisiae* and *Schizosaccharomyces pombe*. FEMS Yeast Res 6, 120–128.

Ariotti N, Murphy S, Hamilton N, Wu L, Green K, Schieber NL, Li P, Martin S, Parton RG (2012). Postlipolytic insulin-dependent remodeling of micro lipid droplets in adipocytes. Mol Biol Cell 23,1826–1837.

Ashe MP, De Long SK, Sachs AB (2000). Glucose depletion rapidly inhibits translation initiation in yeast. Mol Biol Cell 11, 833–848.

Barry RM, Bitbol AF, Lorestani A, Charles EJ, Habrian CH, Hansen JM, Li HJ, Baldwin EP, Wingreen NS, Kollmanand JM, Gitai Z (2014). Large-scale filament formation inhibits the activity of CTP synthetase. eLife 3, e03638.

Bertin A, Nogales E (2016). Characterization of septin ultrastructure in budding yeast using electron tomography. In: Yeast Cytokinesis: Methods and Protocols, eds. A Sanchez-Diaz, P Perez, New York: Springer, 113–123.

Blomberg A, Adler L (1992). Physiology of osmotolerance in fungi. Adv Microb Physiol 33, 145–212.

Bozaquel-Morais BL, Madeira JB, Maya-Monteiro CM, Masuda CA, Montero-Lomeli M (2010). A new fluorescence-based method identifies protein phosphatases regulating lipid droplet metabolism. PLoS One 5, e13692.

Campbell SG, Hoyle NP, Ashe MP (2005). Dynamic cycling of eIF2 through a large eIF2B-containing cytoplasmic body implications for translation control. J Cell Biol 170, 925–934.

Choder M (1993). A growth rate-limiting process in the last growth phase of the yeast life cycle involves RPB4, a subunit of RNA polymerase. J Bacteriol 175, 6358–6363.

Costantini LM, Fossati M, Francolini M, Snapp EL (2012). Assessing the Tendency of Fluorescent Proteins to Oligomerize under Physiologic Conditions. Traffic 13, 643–649.

Desfougères Y, Neumann H, Mayer A (2016). Organelle size control – increasing vacuole content activates SNAREs to augment organelle volume through homotypic fusion. J Cell Sci 129, 2817–2828.

Duncan R, Hershey JW (1983). Identification and quantitation of levels of protein synthesis initiation factors in crude HeLa cell lysates by two-dimensional polyacrylamide gel electrophoresis. J Biol Chem. 258, 7228–7235.

Dupont S, Beney L, Ritt JF, Lherminier J, Gervais P (2010). Lateral reorganization of plasma membrane is involved in the yeast resistance to severe dehydration. Biochim Biophys Acta 1798, 975–985.

Egbe NE, Paget GM, Wang H, Ashe MP (2015). Alcohols inhibit translation to regulate morphogenesis in *C. albicans*. Fungal Genet Biol 77, 50–60.

Frank J (2006). Three-Dimensional Electron Microscopy of Macromolecular Assemblies: Visualization of Biological Molecules in their Native State, Oxford: Oxford University Press.

Gervais P, Marechal PA (1994). Yeast resistance to high levels of osmotic pressure: influence of kinetics. J Food Eng 22, 399–407.

Gordiyenko Y, Schmidt C, Jennings MD, Matak-Vinkovic D, Pavitt GD, Robinson CV (2014). eIF2B is a decameric guanine nucleotide exchange factor with a γ2ε2 tetrameric core. Nat Commun 5, 3902.

Gordiyenko Y, Llácer JL, Ramakrishnan V (2019). Structural basis for the inhibition of translation through eIF2α phosphorylation. Nat Commun 10, 2640.

Gray JV, Petsko GA, Johnston GC, Ringe D, Singer RA, Werner-Washburne M (2004). “Sleeping Beauty”: Quiescence in *Saccharomyces cerevisiae*. Microbiol Mol Biol Rev 68, 187–206.

Gomez E, Pavitt GD (2000). Identification of Domains and Residues within the ε Subunit of Eukaryotic Translation Initiation Factor 2B (eIF2Bε) Required for Guanine Nucleotide Exchange Reveals a Novel Activation Function Promoted by eIF2B Complex Formation. Mol Cell Biol 11, 3965–3976.

Hashimoto T, Segawa H, Okuno M, Kano H, Hamaguchi HO, Haraguchi T, Hiraoka Y, Hasui S, Yamaguchi T, Hirose F, Osumi T (2012). Active involvement of micro-lipid droplets and lipid-droplet-associated proteins in hormone-stimulated lipolysis in adipocytes. J Cell Sci 125, 6127–6136.

Heumann J, Hoenger A, Mastronarde D (2011). Clustering and variance maps for cryo-electron tomography using wedge-masked differences. J Struct Biol 175, 288–299.

Hinnebusch AG, Lorsch JR (2012). The mechanism of eukaryotic translation initiation: new insights and challenges. Cold Spring Harb Perspect Biol 4, a011544.

Hodgson RE, Varanda BA, Ashe MP, Allen KE, Campbell CG (2019). Cellular eIF2B subunit localization: implications for the integrated stress response and its control by small molecule drugs. Mol Biol Cell 30, 942–958.

Ireland LS, Johnston GC, Drebot MA, Dhillon N, DeMaggio AJ, Hoekstra MF, Singer RA (1994). A member of a novel family of yeast ‘Zn-finger’ proteins mediates the transition from stationary phase to cell proliferation. EMBO J 13, 3812–3821.

Jennings MD, Pavitt GD (2014). A new function and complexity for protein translation initiation factor eIF2B. Cell cycle 13, 2660–2665.

Jin M, Klionsky DJ (2014). Regulation of autophagy: modulation of the size and number of autophagosomes. FEBS Lett 588, 2457–2463.

Joyner RP, Tang JH, Helenius J, Dultz E, Brune C, Holt LJ, Huet S, Müller DJ, Weis K (2016). A glucose-starvation response regulates the diffusion of macromolecules. eLife 5, e09376.

Kashiwagi K, Takahashi M, Nishimoto M, Hiyama TB, Higo T, Umehara T, Sakamoto K, Ito T, Yokoyama S (2016). Crystal structure of eukaryotic translation initiation factor 2B. Nature 531, 122–125.

Kukulski W, Schorb M, Welsch S, Picco A, Kaksonen M. Briggs JA (2011). Correlated fluorescence and 3D electron microscopy with high sensitivity and spatial precision. J Cell Biol 192, 111–119.

Kurat CF, Natter K, Petschnigg J, Wolinski H, Scheuringer K, Scholz H, Zimmermann R, Leber R, Zechner R, Kohlwein SD (2006). Obese yeast: triglyceride lipolysis is functionally conserved from mammals to yeast. J Biol Chem 281, 491–500.

Liu JL (2010). Intracellular compartmentation of CTP synthase in *Drosophila*. J Genet Genom 37, 281–296.

Luengo I, Darrow M, Spink M, Sun Y, Dai W, He C, Chiu W, Pridmore T, Ashton A, Duke E, Basham M, French A (2017). SuRVoS: Super-Region Volume Segmentation workbench. J Struct Biol 198, 43–53.

Madeira JB, Masuda CA, Maya-Monteiro CM, Matos GS, Montero-Lomelí M, Bozaquel-Morais BL (2015). TORC1 inhibition induces lipid droplet replenishment in yeast. Mol Cell Biol 35, 737–746.

Martinez de Maranon I, Marechal PA, Gervais P (1996). Passive response of *Saccharomyces cerevisiae* to osmotic shifts: cell volume variations depending on the physiological state. Biochem Biophys Res Commun 227, 519–523.

Mastronarde D (1997). Dual-axis tomography: an approach with alignment methods that preserve resolution. J Struct Biol 120, 343–352.

Mastronarde D (2005). Automated electron microscope tomography using robust prediction of specimen movements. J Struct Biol 152, 36–51.

Meaden PG, Arneborg N, Guldfeldt LU, Siegumfeldt H, Jakobsen M (1999). Endocytosis and vacuolar morphology in *Saccharomyces cerevisiae* are altered in response to ethanol stress or heat shock. Yeast 15, 1211–1222.

Miermont A, Waharte F, Hu S, McClean MN, Bottani S, Léon S, Hersen P (2013). Severe osmotic compression triggers a slowdown of intracellular signaling, which can be explained by molecular crowding. Proc Natl Acad Sci USA 110, 5725–5730.

Morris GJ, Winters L, Coulson GE, Clarke KJ (1986). Effect of osmotic stress on the ultrastructure and viability of the yeast *Saccharomyces cerevisiae*. J Gen Microbiol 132, 2023–2034.

Mourão MA, Hakim JB, Schnell S (2014). Connecting the dots: the effects of macromolecular crowding on cell physiology. Biophys J 107, 2761–2766.

Munder MC, Midtvedt D, Franzmann T, Nüske E, Otto O, Herbig M, Ulbricht E, Müller P, Taubenberger A, Maharana S, Malinovska L, Richter D, Guck J, Zaburdaev V, Alberti S (2016). A pH-driven transition of the cytoplasm from a fluid- to a solid-like state promotes entry into dormancy. eLife 5, e09347.

Munna MS, Humayun S, Noor R (2015). Influence of heat shock and osmotic stresses on the growth and viability of *Saccharomyces cerevisiae* SUBSC01. BMC Research Notes 8, 369.

Narayanaswamy R, Levy M, Tsechansky M, Stovall GM, O’Connell JD, Mirrielees J, Ellington AD, Marcotte EM (2009). Widespread reorganization of metabolic enzymes into reversible assemblies upon nutrient starvation. Proc Natl Acad Sci USA 106, 10147–10152.

Noda T, Ohsumi Y (1998). Tor, a phosphatidylinositol kinase homologue, controls autophagy in yeast. J Biol Chem 273, 3963–3966.

Noree C, Sato BK, Broyer RM, Wilhelm JE (2010). Identification of novel filament-forming proteins in *Saccharomyces cerevisiae* and *Drosophila melanogaster*. J Cell Biol 190, 541–551.

Nüske E, Marini G, Richter D, Leng W, Bogdanova A, Franzmann T, Pigino G, Alberti S (2018). Filament formation by the translation factor eIF2B regulates protein synthesis in starved cells. bioRxiv 467829.

Pavitt GD, Ramaiah KV, Kimball SR, Hinnebusch AG (1998). eIF2 independently binds two distinct eIF2B subcomplexes that catalyze and regulate guanine-nucleotide exchange. Genes dev 12, 514–526.

Petrovska I, Nüske E, Kulasegaran G, Gibson K, Munder MC, Malinovska L, Richter D, Verbavatz JM, Alberti S (2014). Filament formation by metabolic enzymes is a specific adaptation to the energy-depleted cellular state. eLife 3, e02409.

Pettersen EF, Goddard TD, Huang CC, Couch GS, Greenblatt DM, Meng EC, Ferrin TE (2004). UCSF Chimera – a visualization system for exploratory research and analysis. J Comput Chem 25, 1605–1612.

Prouteau M, Desfosses A, Sieben C, Bourgoint C, Mozaffari LN, Demurtas D, Mitra AK, Guichard P, Manley S, Loewith R (2017). TORC1 organized in inhibited domains (TOROIDs) regulate TORC1 activity. Nature 550, 265–269.

Rabouille C, Alberti S (2017). Cell adaptation upon stress: the emerging role of membrane-less compartments. Curr Opin Cell Biol 47, 34–42.

Riback JA, Katanski CD, Kear-Scott JL, Pilipenko EV, Rojek AE, Sosnick TR, Drummond DA (2017). Stress-triggered phase separation is an adaptive, evolutionarily tuned response. Cell 168, 1028–1040.

Rivas G, Fernández JA, Minton AP (2001). Direct observation of the enhancement of noncooperative protein self-assembly by macromolecular crowding: indefinite linear self-association of bacterial cell division protein FtsZ. Proc Natl Acad Sci USA 98, 3150–3155.

Rußmayer H, Buchetics M, Gruber C, Valli M, Grillitsch K, Modarres G, Guerrasio R, Klavins K, Neubauer S, Drexler H, Steiger M, Troyer C, Chalabi AA, Krebiehl G, Sonntag D, Zellnig G, Daum G, Graf AB, Altmann F, Koellensperger G, Hann S, Sauer M, Mattanovich D, Gasser B (2015). Systems-level organization of yeast methylotrophic lifestyle. BMC Biol 13, 80.

Schindelin J, Arganda-Carreras I, Frise E, Kaynig V, Longair M, Pietzsch T, Preibisch S, Rueden C, Saalfeld S, Schmid B, Tinevez JY, White DJ, Hartenstein V, Eliceiri K, Tomancak P, Cardona A (2012). Fiji: an open-source platform for biological-image analysis. Nat Meth 9, 676–682.

Serrano R (1977). Energy requirements for maltose transport in yeast. Eur J Biochem 80, 97–102.

Simonin H, Beney L, Gervais P (2007). Sequence of occurring damages in yeast plasma membrane during dehydration and rehydration: mechanisms of cell death. Biochim Biophys Acta 1768, 1600–1610.

Slaninova I, Sestak S, Svoboda A, Farkas V (2000). Cell wall and cytoskeleton reorganization as the response to hyperosmotic shock in *Saccharomyces cerevisiae*. Arch Microbiol 173, 245–252.

Snapp EL, Hegde RS, Francolini M, Lombardo F, Colombo S, Pedrazzini E, Borgese N, Lippincott-Schwartz J (2003). Formation of stacked ER cisternae by low affinity protein interactions. J Cell Biol 163, 257–269.

Taylor EJ, Campbell S, Griffiths CD, Reid P, Slaven JW, Harrison R, Sims P, Pavitt GD, Delneri D, Ashe MP (2010). Fusel alcohols regulate translation initiation by inhibiting eIF2B to reduce ternary complex in a mechanism that may involve altering the integrity and dynamics of the eIF2B body. Mol Biol Cell 21, 2202–2216.

Thiam AR, Beller M (2017). The why, when and how of lipid droplet diversity. J Cell Sci 130, 315–324.

Thiery JP, Macaya G, Bernardi G (1976). An analysis of eukaryotic genomes by density gradient centrifugation. J Mol Biol 108, 219–235.

Trappe V, Prasad V, Cipelletti L, Segre PN, Weitz DA (2001). Jamming phase diagram for attractive particles. Nature 411, 772–775.

Warner JR (1999). The economics of ribosome biosynthesis in yeast. Trends Biochem Sci 24, 437–440.

Werner-Washburne M, Braun E, Johnston GC, Singer RA (1993). Stationary phase in the yeast *Saccharomyces cerevisiae*. Microbiol Rev 57, 383–401.

Williams DD, Pavitt GD, Proud CG (2001). Characterization of the initiation factor eIF2B and its regulation in *Drosophila melanogaster*. J Biol Chem 276, 3733–3742.

Winderickx J, Holsbeeks I, Lagatie O, Giots F, Thevelein J, de Winde H (2003). From feast to famine; adaptation to nutrient availability in yeast. In: Yeast Stress Responses, eds. S Hohmann, PWH Mager, Berlin, Heidelberg: Springer, 305–386.

Woodruff JB, Ferreira Gomes B, Widlund PO, Mahamid J, Honigmann A, Hyman AA (2017). The Centrosome Is a Selective Condensate that Nucleates Microtubules by Concentrating Tubulin. Cell 169, 1066–1077.

Zacharias DA, Violin JD, Newton AC, Tsien RY (2002). Partitioning of lipid-modified monomeric GFPs into membrane microdomains of live cells. Science 296, 913–916.

Zhou HX, Rivas G, Minton AP (2008). Macromolecular crowding and confinement: biochemical, biophysical, and potential physiological consequences. Annu Rev Biophys 37, 375–397.

